# Th9-endothelial cell crosstalk promotes inflammatory atherosclerotic cardiovascular disease

**DOI:** 10.1101/2025.05.05.652112

**Authors:** Ishita Baral, Yvonne Baumer, Aarohan Burma, McKella Sylvester, Kyle Jones, Moses Mwesigwa Kitakule, Amit Dey, Justin Rodante, Cristhian A. Gutierrez-Huerta, Bhavani Taramangalam, Julio Diaz Perez, Liang Guo, Alyssa Grogan, Tatsuya Shiraki, Aloke Finn, Aran Son, Kyoungin Cho, Jaspal Khilliian, Marcus Chen, Tiffany M. Powell-Wiley, Pamela A. Frischmeyer-Guerrerio, Joshua D. Milner, Nehal N Mehta, Guido H. Falduto, Daniella M. Schwartz

## Abstract

Atherosclerotic cardiovascular disease (ASCVD) is a leading cause of death, and understanding its pathogenic drivers is critical for effective prevention and treatment. Inflammation has a critical role in ASCVD, and patients with inflammatory diseases are at increased risk. However, the key inflammatory mediator promoting ASCVD are incompletely understood, a major barrier when targeting inflammation to prevent ASCVD. Here, we found that interleukin-9 (IL-9) producing T helper cells (Th9) were significantly associated with ASCVD in patients with the autoimmune disease psoriasis. Th9 cells were poised to migrate to coronary vessels and were identified in atherosclerotic plaque. *In vivo*, murine inflammatory atherogenesis was prevented by IL-9 blockade and by IL-9 receptor (IL-9R) deletion in endothelial cells. In human arterial endothelial cells, IL-9R/STAT3 signaling promoted endothelial dysfunction, angiogenesis, and release of leukocyte chemoattractants. These findings suggest that in autoimmune diseases like psoriasis, Th9/IL-9 promote atherosclerosis by directly targeting endothelial cells, and that IL-9R/STAT3 signaling could be a promising therapeutic target for ASCVD.

**eTOC Summary:** Baral et. al. investigate individuals who are at high risk for atherosclerosis due to underlying inflammatory disease and use mouse models of cardiovascular disease to uncover a role for interleukin-9-producing T helper 9 cells in the pathogenesis of inflammatory atherosclerosis.

## Introduction

Atherosclerotic cardiovascular disease (ASCVD) is the leading cause of death worldwide, accounting for more than 10 million deaths per year (Vaduganathan et al., 2022). Understanding the pathogenic drivers of atherosclerotic disease is critical for effective primary and secondary prevention. The contribution of inflammation to ASCVD has been recognized for decades (Libby et al., 2002). Atherogenesis, or the formation of arterial plaque, involves endothelial cell dysfunction, leading to recruitment of immune cells like monocytes and T cells (Libby et al., 2002; Schwartz et al., 2020). These cells infiltrate lipid-rich plaque and react to local mediators by releasing proinflammatory cytokines that promote plaque core necrosis, recruitment of additional inflammatory cells, and worsening endothelial cell dysfunction (Libby et al., 2002; Wolf and Ley, 2019).

Major efforts have been made to identify pathogenic drivers of inflammatory CVD that could be targeted for the therapeutic benefit of at-risk individuals (Libby et al., 2002; Wolf and Ley, 2019). Nonetheless, current strategies target metabolic factors like dyslipidemia and glucose intolerance (Vaduganathan et al., 2022). These strategies may indirectly reduce inflammatory CVD through metabolic-inflammatory interactions (Wolf and Ley, 2019) but do not directly target inflammatory mediators. Multiple large clinical trials have investigated the efficacy of broad-spectrum and targeted anti-inflammatory therapies in CVD prevention (Mitsis et al., 2024; Ridker et al., 2017; Tardif et al., 2019). However, results for many of these agents have been disappointing, while other promising immunomodulatory agents have been judged too high-risk for widespread clinical use. This may be because the inflammatory mediators promoting ASCVD are diverse and can differ between patients and cohorts (Doring et al., 2024). This heterogeneity can present significant challenges to identifying and stratifying the individuals most likely to benefit from targeted immunomodulatory therapies. Indeed, only one broad-spectrum medication, colchicine, is clinically used to target inflammation in CVD patients (Mitsis et al., 2024; Ridker et al., 2017; Tardif et al., 2019). Nonetheless, the strong evidence implicating inflammation in the pathogenesis of ASCVD suggests that improved patient selection and identification of key disease-causal cytokines and immune cells can help to better stratify patients and identify novel therapeutic targets (Doring et al., 2024; Wolf and Ley, 2019).

The relationship between inflammation and atherogenesis is most evident in patients with systemic autoimmune diseases (Conrad et al., 2022; Manning et al., 2025). Not only are these patients at significantly increased risk of ASCVD (Conrad et al., 2022; Manning et al., 2025), but they undergo treatment with cytokine-targeting agents that can also modulate their CVD risk (Fragoulis et al., 2021). Amongst these systemic autoimmune diseases, psoriasis stands out as a particularly useful “human model” of chronic inflammatory atherogenesis (Harrington et al., 2017). Psoriasis is a systemic disease that commonly affects the skin and joints, and can also affect the eyes, gastrointestinal tract, and bones (Griffiths et al., 2021). Psoriatic disease severity has a dose-response relationship with cardiometabolic dysregulation, adipose inflammation, and ASCVD (Eder et al., 2015; Harrington et al., 2017; Schwartz et al., 2022). Mechanistically, psoriasis has been linked to pro-atherogenic pathways including immune activation, metabolic dysregulation, lipoprotein abnormalities, and endothelial dysfunction (Baumer et al., 2017; Griffiths et al., 2021; Harrington et al., 2017; Visser et al., 2021). However, the specific pathogenic drivers and mechanisms underlying inflammatory CVD in patients with psoriasis remain incompletely understood. While immunomodulatory treatments are thought to reduce psoriatic CVD risk in some patients, results of real-world data and clinical trials have been contradictory (Chen et al., 2024; Ding et al., 2024; Galajda et al., 2024). This could be related to protective roles for some cytokines in stabilizing arterial plaque (Madhur et al., 2011; Mallat et al., 1999), heterogeneity between patient populations (Chen et al., 2024; Ding et al., 2024), or failure to stratify patients according to disease-associated biomarkers prior to analysis or treatment. These observations suggest that other pathogenic cytokines and immune cells may be the key drivers of inflammatory atherogenesis and potential therapeutic targets.

Here, we found that psoriatic CVD was significantly associated with expansion of circulating interleukin-9 (IL-9) producing CD4^+^ T cells. These IL-9-producing T cells expressed a T helper 9 (Th9) lineage-defining transcriptomic signature, were poised to migrate to arterial plaque, and were found in murine and human atherosclerotic plaques. *In vivo*, germline and endothelial IL-9 receptor deletion ameliorated murine inflammatory atherogenesis, as did therapeutic IL-9 blockade. *In vitro*, IL-9 directly targeted arterial endothelial cells to induce endothelial dysfunction and promote leukocyte migration. Our findings implicate Th9 cells and IL-9 in the pathogenesis of inflammatory atherogenesis, providing a novel link that could represent an attractive therapeutic target to prevent and treat CVD in patients with autoimmune and inflammatory diseases.

## Results

### Psoriasis is associated with expansion of circulating Th9 cells

Based on prior reports that psoriasis is an IL-9/Th9-high state (Clark and Schlapbach, 2017; Micosse et al., 2019; Schlapbach et al., 2014), we first phenotyped circulating T helper subsets from psoriatic (n = 36) and non-psoriatic subjects (n = 47). As a positive comparator, we profiled circulating Th17 cells (**Fig. 1a-b**) in psoriatic and non-psoriatic subjects, measuring the percentage of memory (CD4^+^CD45RO^+^) Th cells that expressed IL-17A. As expected, IL-17A^+^ memory T cells were significantly expanded in subjects with psoriatic disease (**Fig 1b**). Most subjects had mild to moderate disease, and accordingly the percentage of circulating Th17 cells was only modestly elevated. We did not find expansion of Th1 cells (IFN-γ^+^), Th2 cells (IL-4^+^, IL-13^+^, or IL-5^+^), or of memory CD4^+^ T cells expressing IL-2, IL-10, IL-21, or TNF-α (**Fig S1a-h**). However, circulating IL-9^+^ memory CD4+ T cells were also significantly expanded in psoriatic subjects (**Fig 1c-d**), indicating that psoriasis is a systemic Th9-high state.

**Fig 1.**
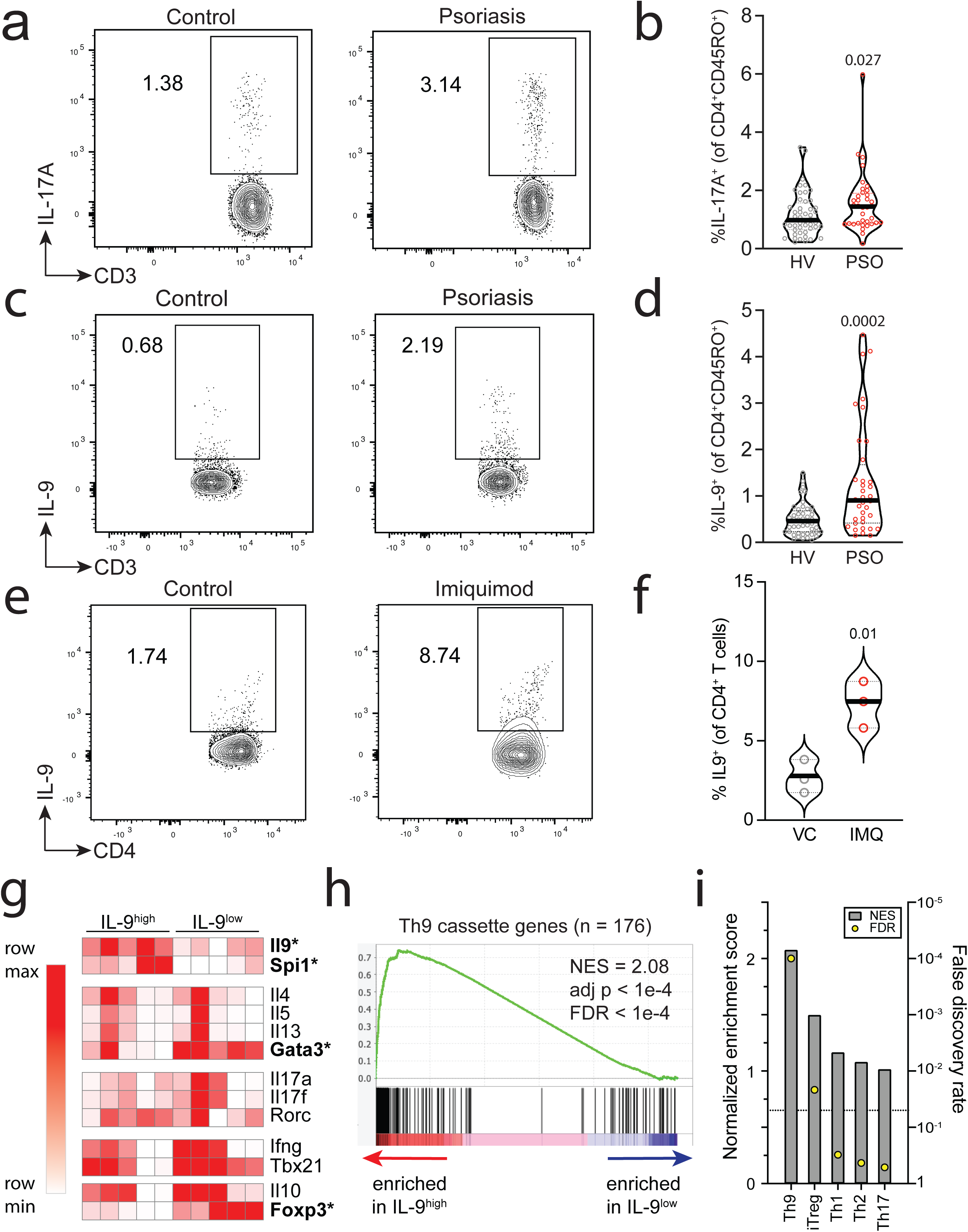
a,b. Flow (a) and violin (b) plot show circulating memory Th17 (CD45^+^/CD3^+^/CD4^+^/CD45RO^+^/IL17A^+^) cells in non-psoriatic healthy volunteers (HV, n = 47, grey) and patients with psoriasis (PSO, n = 36, red). c,d. Flow (c) and violin (d) plot show circulating memory Th9 (CD45+/CD3+/CD4+/CD45RO+/IL9+) cells in non-psoriatic healthy volunteers (HV, n = 47, grey) and patients with psoriasis (PSO, n = 36, red). p-values, Mann-Whitney. e,f. Flow (e) and violin (f) plot show splenic Th9 (CD45+/TCR-β+/ CD4+/IL9+) cells in mice treated with three rounds of imiquimod (IMQ) or vehicle (**Fig S1j**) p-values, unpaired t-test, n = 3 g. Heatmap shows expression (normalized across each row) of lineage-defining transcription factors and effector cytokines for Th9, Th2, Th17, Th1, and Treg subsets. * adjusted p < 0.05, DESeq2. h. GSEA plot shows enrichment of “allergic Th9 cassette” in splenic GFP+ (Th9) vs. GFP-(non-Th9) cells from INFER (IL-9 reporter) mice treated with IMQ. i. Bar graphs show normalized enrichment scores and false discovery rate for enrichment (GSEA) of lineage defining cassettes for the Th9, iTreg (induced regulatory T cell), Th1, Th2, and Th17 subsets in splenic GFP+ (Th9) and GFP-(non-Th9) cells from INFER mice treated with IMQ.

In subjects with allergic disease, IL-9^+^ T helper cells have been described as an early or inflammatory subset of Th2 cells (Alladina et al., 2023; Micosse et al., 2019). In most subjects with psoriasis, Th9 cells did not co-express Th2 cytokines (IL-4, IL-13, IL-5) or IL-17A but were instead single positive cells (**Fig S1i**). To more comprehensively investigate the identity of Th9 cells in the context of psoriatic disease, we utilized IL-9 reporter (INFER) mice (Licona-Limon et al., 2013), exposing the mice to multiple topical applications of the toll-like receptor (TLR) -7/9 agonist imiquimod (IMQ) to induce chronic/recurrent psoriasiform inflammation (**Fig S1j**) (van der Fits et al., 2009). IMQ induced expansion of circulating Th9 cells expressing GFP/IL-9, confirming that this model could be used to study Th9 cells in the context of inflammatory/autoimmune skin disease (**Fig 1e**-**f**). Analysis of the transcriptomes of circulating IL-9^+^ CD4^+^ T cells revealed significant (adj p < 0.05, DESeq2) differential upregulation of *Il9* and of the Th9 lineage-defining transcription factor (TF) *Spi1* (**Fig 1g**). We had previously identified a cassette of Th9 lineage-defining genes(Son et al., 2023), which was also significantly enriched (FDR < 0.05, GeneSet Enrichment Analysis, GSEA) in circulating IL-9^+^ T helper cells from IMQ-treated mice (**Fig 1h**). Th9 cells from IMQ-treated mice did not exhibit differential upregulation of other T helper lineage-defining effector cytokines, but they did have increased expression of the Th2 master TF *Gata3* and the regulatory T cell (Treg) TF *Foxp3* (**Fig 1g**). Analysis of lineage-specific cassettes for other T helper subsets (Schwartz et al., 2019) revealed weak enrichment for Treg-specific genes and no enrichment for other T helper subsets (**Fig 1i**). Together, these results confirmed that murine psoriasis models recapitulated the systemic Th9-high state seen in human psoriatic disease, and that psoriasis-associated Th9 cells express similar lineage-defining genes to allergy-associated Th9 cells.

### Th9-high state is associated with coronary artery disease in psoriatic subjects

Th9 cells and IL-9 have previously been linked to skin inflammation in patients with psoriasis and atopic dermatitis (Clark and Schlapbach, 2017). We therefore hypothesized that Th9-high state might correlate with psoriatic skin disease severity. Psoriasis Area Severity Scores (PASI) had been calculated for a subset of patients (n = 20) but did not correlate with Th9 expansion in peripheral blood (**Supplementary Table 1**) This led us to question whether Th9 cells might promote disease at other sites. Patients with psoriasis are at increased risk of psoriatic arthritis, inflammatory bowel disease, and cardiovascular disease, all of which have been associated with Th9 cells in other contexts (Ciccia et al., 2016; Gerlach et al., 2014; Rauber et al., 2017; Son et al., 2024; Zhang et al., 2015). A group of 19 psoriasis patients had undergone cardiovascular disease screening with coronary artery computed tomography angiography (CCTA) (**Fig S1k**). We analyzed CCTA to calculate non-calcified coronary plaque burden, a high-risk type of coronary plaque that is associated with increased mortality and myocardial infarction risk (Abdelrahman et al., 2020). Th9-high state was significantly associated with both total and non-calcified burden; this association remained significant after adjusting for traditional cardiovascular risk factors like body mass index, Framingham risk score, and statin use (**Table 1, Supplementary Table 1**). Similar associations were not seen for other T helper cytokines and subsets (**Supplementary Table 2**). These data strongly suggested that a Th9-high state is associated with developent of atherosclerotic coronary artery disease in psoriasis patients.

**Table 1.** IL-9+ CD45RO+ CD4+ T cells and Non-Calcified Burden Adjusted Regression Analysis.

| Model | Standardized Beta | P-Value* |
| --- | --- | --- |
| Unadjusted | 0.28 | **0.039** |
| Adjusted for BMI, FRS, and Statin Use | 0.23 | **0.002** |

\*Multivariable regression analysis between non-calcified burden and %IL-9+ CD45RO+ CD4+ T cells unadjusted or adjusted for BMI, FRS, and statin use.

### Circulating Th9 cells express a pro-atherogenic program in psoriatic subjects

The strong association of Th9 expansion with non-calcified coronary burden led us to hypothesize that Th9 cells might be poised to infiltrate the vessel wall. Analysis of circulating Th9 cells from IMQ-treated mice revealed 1819 Th9^high^ genes (FC > 2, padj < 0.05, DESeq2, IL-9^high^ vs. IL-9^low^, **Fig S1l**),1330 of which were highly differentially expressed in Th9^high^ cells (FC > 4, padj < 0.05, DESeq2, IL-9^high^ vs. IL-9^low^). These highly differentially expressed genes (DEGs) were enriched for multiple pathways associated with inflammatory atherogenesis including vascular adhesion, angiogenesis, and lipid metabolism (**Fig 2a-c**). Genes associated with lymphocyte homing to the heart were significantly upregulated in circulating Th9 cells (**Fig 2d**): these included *Met*, *Itgam, Itgb2,* and *Cxcr2* (Komarowska et al., 2015; Marchini et al., 2021; Wang et al., 2016). Together, these findings suggested that circulating Th9 cells in psoriatic subjects are poised to traffic to the heart and undergo recruitment to coronary arteries.

**Fig 2.**
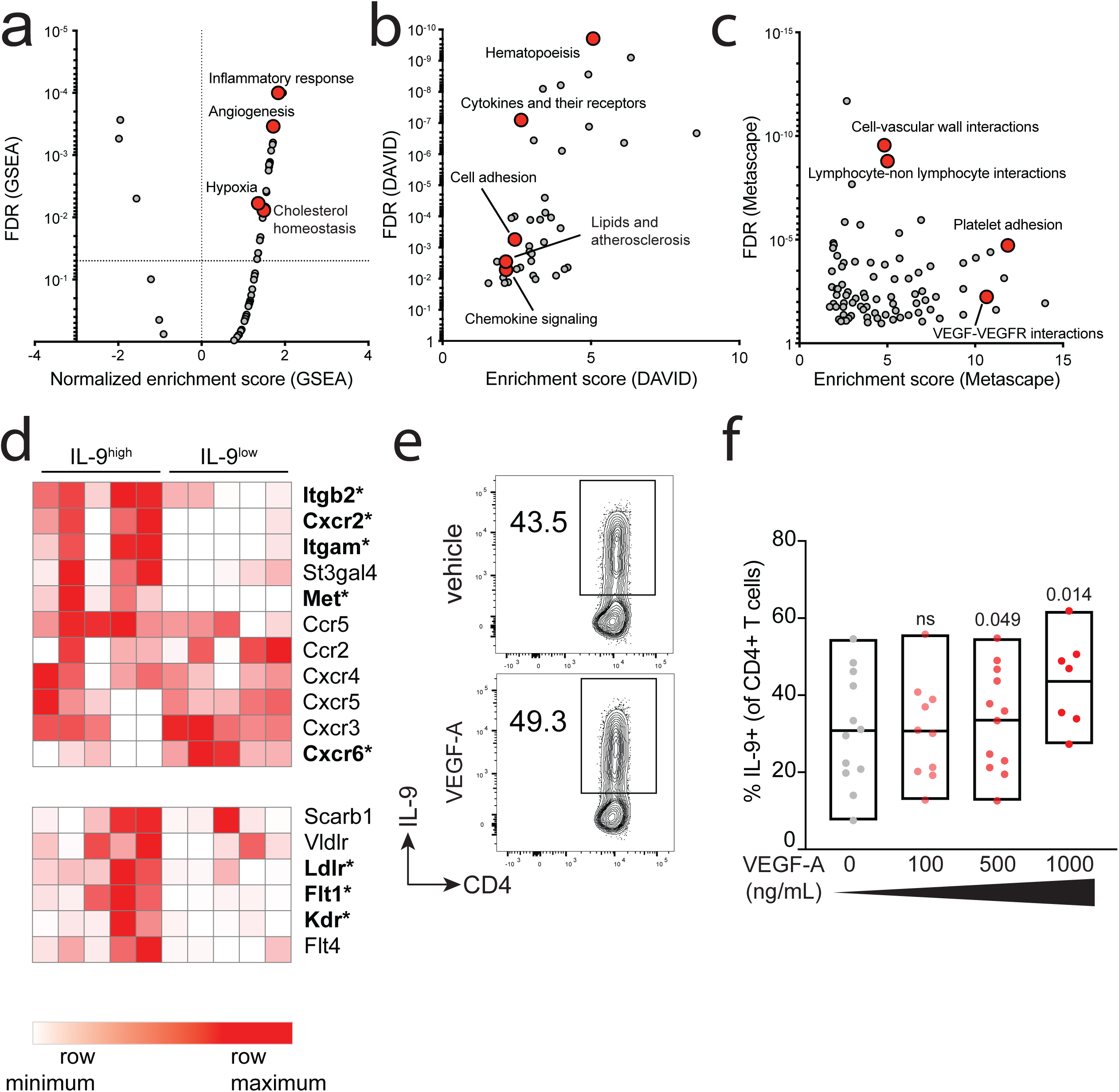
a-c. Scatter plots show enrichment scores vs. false discovery rates calculated using GSEA (a), DAVID (b), or Metascape (c). All enrichments were calculated for splenic GFP+ (Th9) cells compared to GFP-(non-Th9) cells isolated from INFER (IL-9-reporter) mice treated with imiquimod (IMQ) as in **Fig S1j**. d. heatmap shows expression (normalized across each row) of genes important for trafficking to the blood vessel and/or heart (top panel) or genes encoding receptors of pro-angiogenic factors (bottom panel). * adjusted p < 0.05, DESeq2. e,f. Flow plot (e) and bar graph (f) show percent murine *in vitro* differentiated murine Th9 cells (of CD4+ cells) differentiated in the presence of vehicle (grey) or oxidized LDL (oxLDL, red). g,h. Flow plot (g) and bar graph (h) show percent murine *in vitro* differentiated murine Th9 cells (of CD4+ cells) differentiated in the presence of vehicle (grey) or escalating concentrations of VEGF-A (red). p-values use ANOVA with multiple comparison adjustment.

After trafficking to the coronary arteries, CD4^+^ T cells are exposed to pro-atherogenic factors that can enhance the expansion of inflammatory T helper subset and production of proinflammatory cytokines (Saigusa et al., 2020). This led us to hypothesize that Th9 cells might be activated and expand within the pro-atherogenic plaque environment. Among the pathways differentially upregulated in psoriatic Th9 cells, we identified VEGF-VEGFR signaling and lipid metabolism as potential drivers of Th9 differentiation and exspansion within the plaque environment (**Fig 2a-c**). The genes encoding VEGFR-1, VEGFR-2, and LDL receptor were significantly upregulated in Th9 cells relative to non-Th9 cells (**Fig 2d**). We therefore differentiated naïve murine CD4^+^ T cells under Th9-promoting conditions in the presence of VEGF-A and oxidized LDL. Oxidized LDL repressed Th9 differentiation, whereas VEGF-A significantly enhanced Th9 differentiation, implicating VEGF-A as a proatherogenic factor that could enhance Th9 differentiation within arterial plaque (**Fig 2e-f, S1m**).

### Th9 cells infiltrate the atherosclerotic plaque of psoriatic subjects

Our findings that psoriasis-associated Th9 cells expressed coronary artery trafficking markers and were modulated by pro-atherogenic factors led us to hypothesize that atherosclerotic plaques from psoriatic subjects would be characterized by Th9 infiltration. Because T-cell-driven atherogenesis is also a central driver of non-psoriatic cardiovascular disease (Saigusa et al., 2020), we first investigated Th9 cell infiltration in atherosclerotic plaque from non-psoriatic subjects. Using CITE-seq of sorted T cells from the peripheral blood and carotid plaques of patients with atherosclerotic cardiovascular disease, we were able to identify a population of CD4^+^ T cells expressing the Th9 lineage-defining transcription factor *SPI1* (**Fig S2a**) (Chang et al., 2010; Fernandez et al., 2019; Gerlach et al., 2014; Kaplan, 2017). Th9 lineage-defining genes (Son et al., 2023) were significantly enriched (FDR < 0.05, GSEA) in plaque-infiltrating *SPI1*^+^CD4^+^ T cells, but not in circulating *SPI1*^+^CD4^+^ T cells (**Fig 3a, S2b**). Analysis of the 347 genes most closely correlated with *SPI1* expression (Pearson correlation) revealed enrichment for canonical Th9-associated pathways like IL-2 induction, response to IL-4, and response to TGF-β (**Fig 3b**). *SPI1*-associated genes were also enriched for PPAR signaling, which is important for Th9 differentiation, CD4^+^ T cell activation, lipid metabolism, and cardiovascular function (Micosse et al., 2019; Montaigne et al., 2021; Straus and Glass, 2007). Other enriched pathways included cellular adhesion and responses to proatherogenic pathways downstream of hydrogen peroxide (Batty et al., 2022), NOD2 (Liu et al., 2013), oxidized phospholipids (Ox-PL) (Lee et al., 2012), and TLR2 (Mullick et al., 2008). Together, these data suggest that Th9 cells infiltrate atherosclerotic plaque and respond to plaque-resident inflammatory mediators. To specifically investigate Th9 infiltration in coronary plaque from subjects with psoriasis, we obtained postmortem samples from subjects with psoriasis and coronary artery disease. Sampling 6 arteries from 3 deceased subjects with a documented history of psoriatic disease revealed multiple PU.1^+^/CD3^+^ and PU.1^+^/CD4^+^ cells within necrotic lipid-rich coronary artery plaque (**Fig 3c-d**).

**Fig 3.**
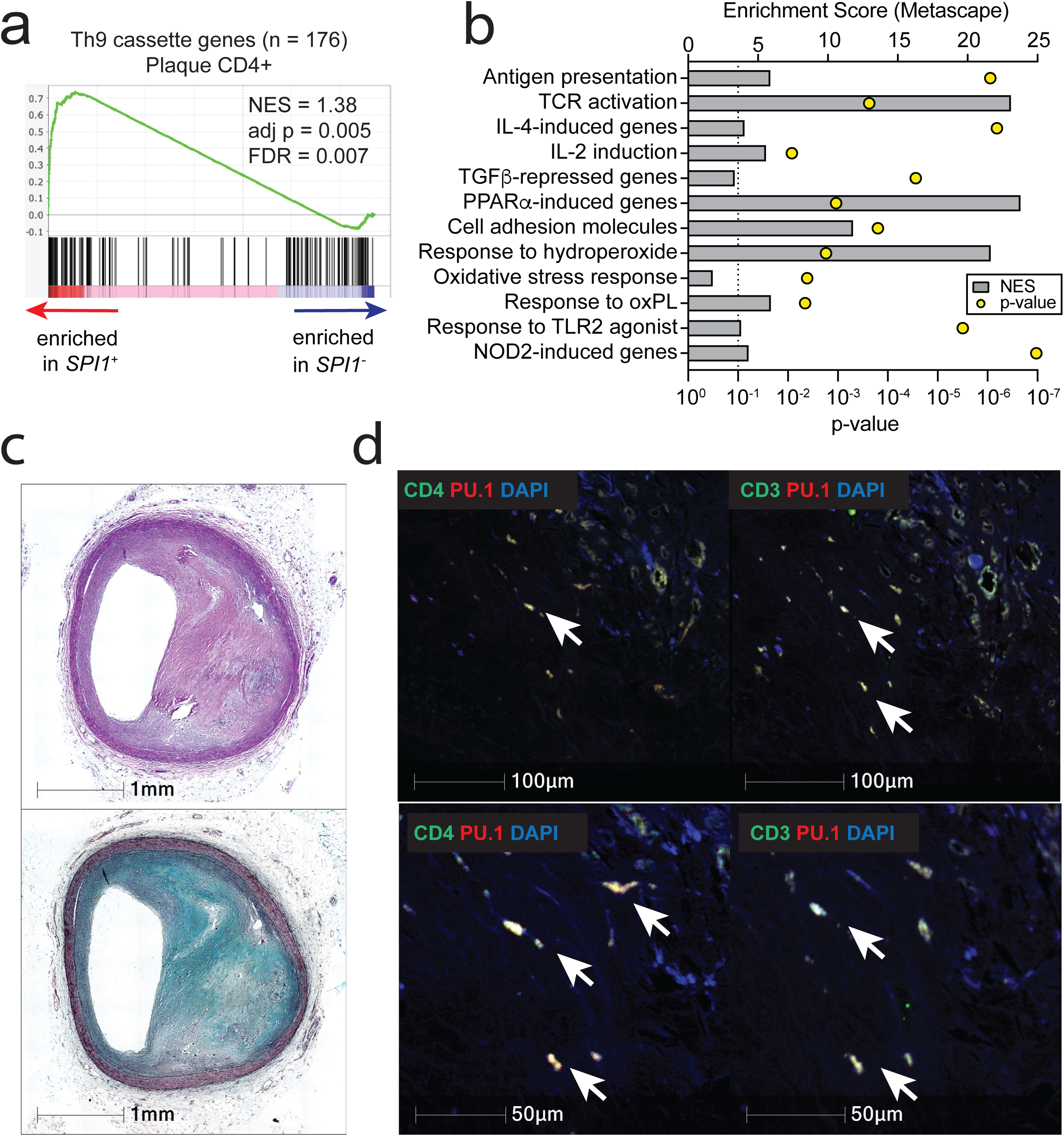
a. GSEA plot shows enrichment of “allergic Th9 cassette” in carotid plaque resident *SPI1*+ (Th9) vs. *SPI1*-(non-Th9) cells from public dataset of plaque-resident CD4+ T cells from humans undergoing carotid endarterectomy. b. Bar graphs show enrichment scores and p-values (Metascape) for pathways enriched in genes whose expression correlates (Pearson≥0.2) to *SPI1* expression in carotid plaque-resident CD4+ T cells. c-e. representative images of coronary artery plaque from subject with psoriasis and atherosclerotic coronary artery disease show H&E and MOVAT staining (c), immunofluorescence staining (d) for the Th9 lineage-defining transcription factor PU.1 (red, encoded by *SPI1*) and CD3 (green), and immunofluorescence staining (e) for PU.1 (red) and CD4 (red).

To dissect the mechanisms by which Th9-derived IL-9 might promote inflammatory atherosclerosis, we next sought to develop a murine model in which skin inflammation was associated with accelerated atherogenesis. We therefore exposed *ApoE^-/-^* mice, which are atherosclerosis-prone mice, to topical IMQ while feeding them a high fat diet (HFD) to induce atherogenesis (**Fig 4a**). Because the relative risk of psoriatic CVD is highest in young patients with severe skin disease (Samarasekera et al., 2013), we used an early time point of 6 weeks HFD. We induced 3 cycles of inflammation over 7-10% body surface area to model moderate-severe psoriasiform skin disease (Geale and Schmitt-Egenolf, 2022). Compared with HFD or IMQ alone, the combination of HFD and IMQ significantly increased lipid-rich plaque in the aortic arch (**Fig 4b**). In mice exposed to HFD, IMQ-treatment nonsignificantly induced CD4^+^ T cell infiltration into skin, while IFN-γ+, IL-13^+^, IL-17A^+^, and IL-9^+^ CD4^+^ T cells were significantly higher in murine skin of IMQ-treated mice (**Fig 4d-e, S2c-d**). IMQ-treatment also significantly induced IL-9^+^ CD4^+^ T cell infiltration into murine aortas but did not induce other T helper subsets (**Fig 4f-g, S4d**). These findings confirmed that skin inflammation is associated with accelerated atherogenesis and increased aortic Th9 cell infiltration.

**Fig 4.**
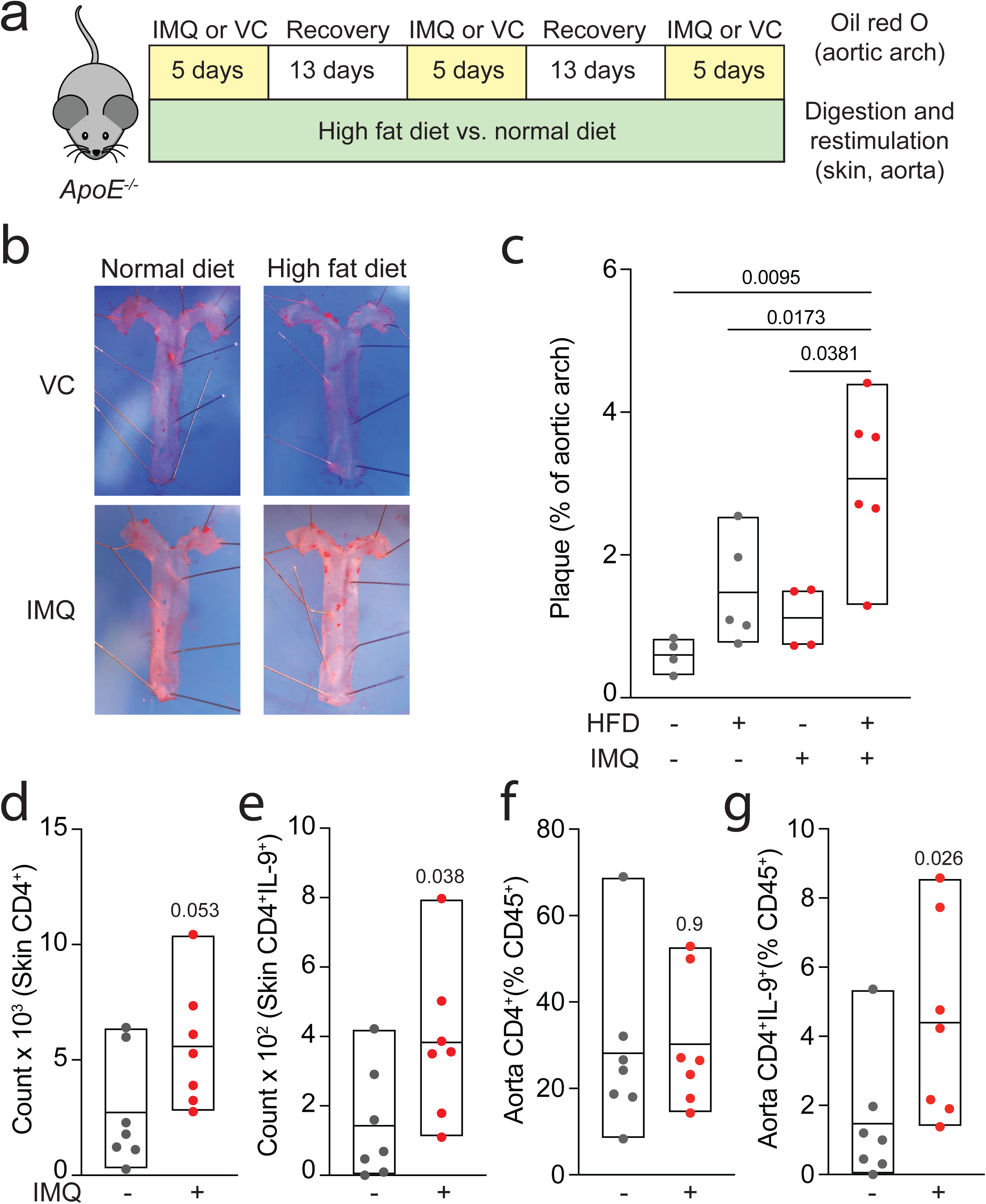
a. Schematic shows design of IMQ-HFD model of psoriatic atherogenesis. *ApoE^-/-^* mice were exposed to three 5-day rounds of imiquimod (IMQ) or vanicream (VC) with two 13-day recovery periods, while being fed high-fat diet (HFD) or normal chow. After 6 weeks, mice were euthanized. The thoracic aorta was stained to visualize lipid-rich plaque (Oil-Red-O), while the skin and abdominal aorta was digested for immune cell analysis, including restimulation for T helper subset profiling (flow cytometry). b,c. Representative images (b) and bar graph (c) show pooled oil red O (+) lipid-rich plaque as percentage of aortic arch in *ApoE^-/-^*mice treated with normal diet or high fat diet while being exposed to VC (grey) or IMQ (red). d,e. Bar graphs show counts of total (d) and IL-9+ (e) skin-infiltrating CD4+ T cells in mice treated with VC (grey) or IMQ (red). f,g. counts of total (d) and IL-9+ (e) aorta-infiltrating CD4+ T cells in mice treated with VC (grey) or IMQ (red). For all experiments: p-values, Mann-Whitney.

### IL-9 promotes inflammatory atherogenesis

Because our murine and human data strongly implicated Th9 cells and IL-9 in psoriatic atherosclerosis, we next hypothesized that IL-9 might directly promote inflammatory atherogenesis. To test this hypothesis, we exposed *Il9r^-/-^ApoE^-/-^* and *ApoE^-/-^* mice to IMQ for 6 weeks while on HFD (**Fig S2e**). *Il9r^-/-^ApoE^-/-^* mice had less skin infiltration of neutrophils, eosinophils, and inflammatory macrophages (**Fig S2f**) compared with *ApoE^-/-^* mice, but there was no significant difference in other immune cell populations (**Fig S2g**). IL-9 receptor deletion also reduced lipid-rich aortic plaque formation (**Fig 5a,b**) and aortic infiltration of neutrophils and inflammatory macrophages (**Fig 5c,d**) but had no significant effect on other immune cell subsets (**S2h**). Like IL-9 receptor-deficient mice, *ApoE^-/-^* mice treated with IL-9-neutralizing antibodies while exposed to IMQ and HFD (**Fig S3a**) were protected from lipid-rich aortic plaque formation (**Fig 5e, f**). However, IL-9-neutralizing antibodies did not significantly reduce aorta-infiltrating (**Fig 5g-h, S3b**) or skin-infiltrating (**Fig S3c**) immune cell populations in IMQ-treated mice.

**Fig 5.**
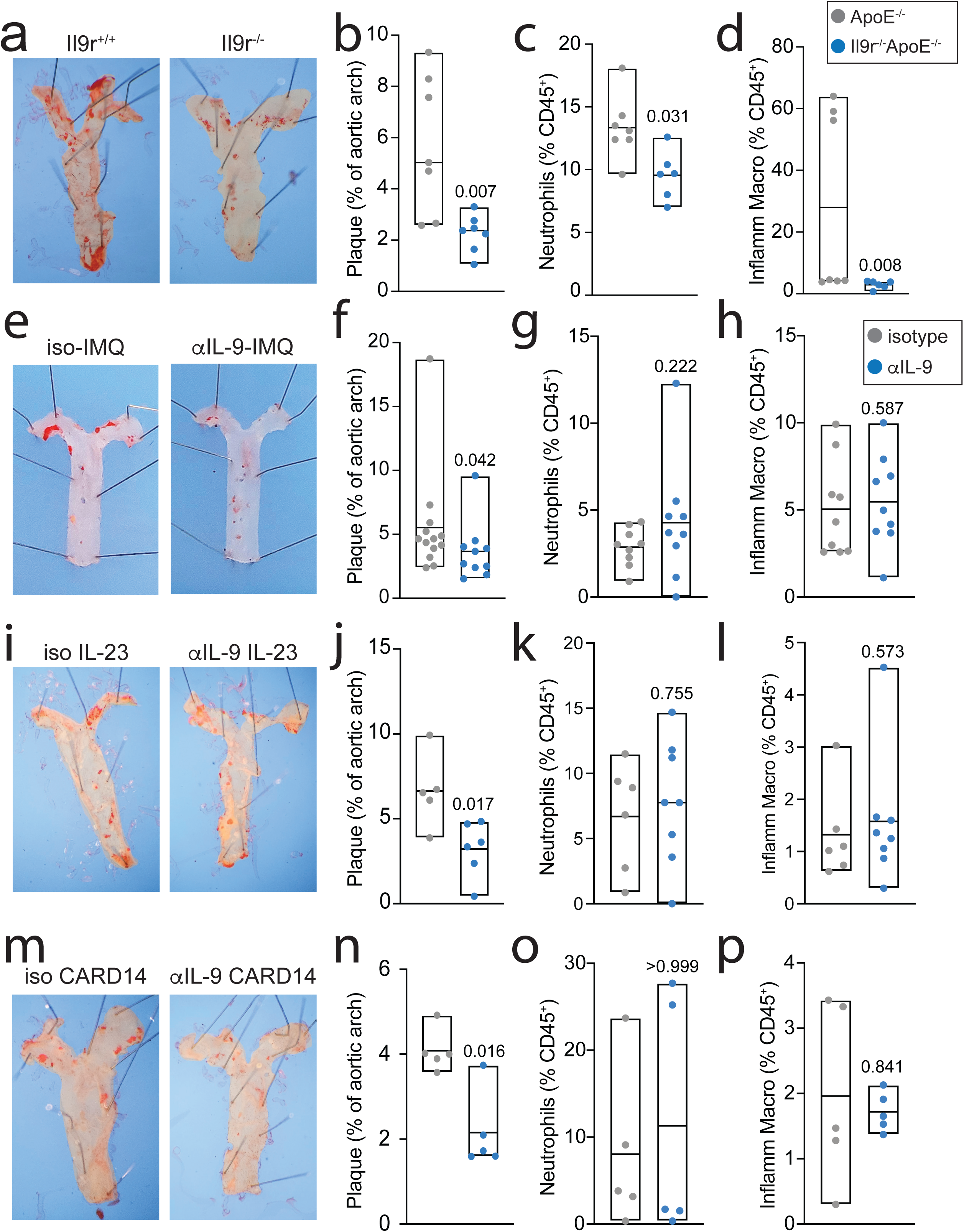
a,b. Representative images (a) and bar graph (b) show pooled oil red O (+) lipid-rich plaque as percentage of aortic arch in *ApoE^-/-^* (grey) and *Il9r^-/-^ApoE^-/-^* mice (blue) exposed to three rounds of imiquimod (IMQ, Fig 4a**, S2j**) while being exposed to high-fat diet (HFD). c,d. Bar graphs show percent neutrophils (c) and Ly6C+ inflammatory macrophages (d) of aorta-infiltrating CD45^+^ cells in *ApoE^-/-^* (grey) and *Il9r^-/-^ApoE^-/-^*mice (blue). e,f. Representative images (e) and bar graph (f) show pooled oil red O (+) lipid-rich plaque as percentage of aortic arch in *ApoE^-/-^* mice exposed to IMQ-HFD while treated with isotype (grey) or αIL-9 (blue). g,h Bar graphs show percent neutrophils (g) and Ly6C+ inflammatory macrophages (h) of aorta-infiltrating CD45^+^ cells in *ApoE^-/-^* mice treated with IMQ-HFD and isotype (grey) or αIL-9 (blue). i,j. Representative images (e) and bar graph (f) show pooled oil red O (+) lipid-rich plaque as percentage of aortic arch in *ApoE^-/-^*mice exposed to three rounds of intradermal IL-23 (**Fig S3a**) while being exposed to HFD and treated with isotype (grey) or αIL-9 (blue). k,l Bar graphs show percent neutrophils (k) and Ly6C+ inflammatory macrophages (l) of aorta-infiltrating CD45^+^ cells in *ApoE^-/-^* mice treated with IL-23-HFD and isotype (grey) or αIL-9 (blue). m,n. Representative images (m) and bar graph (n) show pooled oil red O (+) lipid-rich plaque as percentage of aortic arch in *Card14^ΔE138^ApoE^-/-^* mice (**Fig S3k**) exposed to HFD and treated with isotype (grey) or αIL-9 (blue). o,p Bar graphs show percent neutrophils (o) and Ly6C+ inflammatory macrophages (p) of aorta-infiltrating CD45^+^ cells in *Card14^ΔE138^ApoE^-/-^*mice treated with HFD and isotype (grey) or αIL-9 (blue). For all experiments: p-values, Mann-Whitney.

Because IMQ can induce off-target and unintended consequences that may not fully recapitulate psoriatic inflammation (Gangwar et al., 2022), we next sought to confirm our findings in two other models of psoriasis: intradermal IL-23 injections (**Fig S3a**) and *Card14^ΔE138^* (**Fig S3d**), a gain-of-function (GOF) mutation that is orthologous to and phenocopies human monogenic psoriasis due to *CARD14*-GOF (Gangwar et al., 2022; Wang et al., 2018). In both models, IL-9-neutralizing antibodies reduced lipid-rich aortic plaque formation but failed to significantly reduce plaque-infiltrating or skin-infiltrating immune cell populations, recapitulating the phenotype of IMQ-treated mice (**Fig 5i-p, S3d-g**). These findings indicated that IL-9 promotes early inflammatory atherogenesis and aortic plaque formation, and that these effects might be independent of immune cell recruitment to the atherosclerotic plaque.

### IL9R signaling in endothelial cells promotes inflammatory atherogenesis

Our findings in IL-9R-deficient and αIL-9-treated mice led us to hypothesize that IL-9 might promote inflammatory atherogenesis through direct results on endothelial cells. To test this hypothesis, we generated a murine model in which LoxP sites were inserted to flank exons 2-5 of the *Il9r* gene, allowing us to conditionally delete IL-9 receptor (**Fig 6a**). To confirm that LoxP site insertion did not disrupt endogenous IL-9 receptor signaling, we differentiated in-vitro derived germinal center memory B cells, which phosphorylate STAT3 and STAT5 in response to IL-9 (Takatsuka et al., 2018), from *Il9r^f^*^/f^ and WT mice (**Fig S4a**). WT and *Il9*^f/f^ derived B cells displayed similar responses to treatment with IL-9, confirming that LoxP insertion did not disrupt endogenous IL9 receptor signaling (**Fig S4b**). Treating memory B cells with Tat-Cre *in vitro* significantly reduced IL-9 receptor expression (**Fig S4c**), confirming gene inactivation.

**Fig 6.**
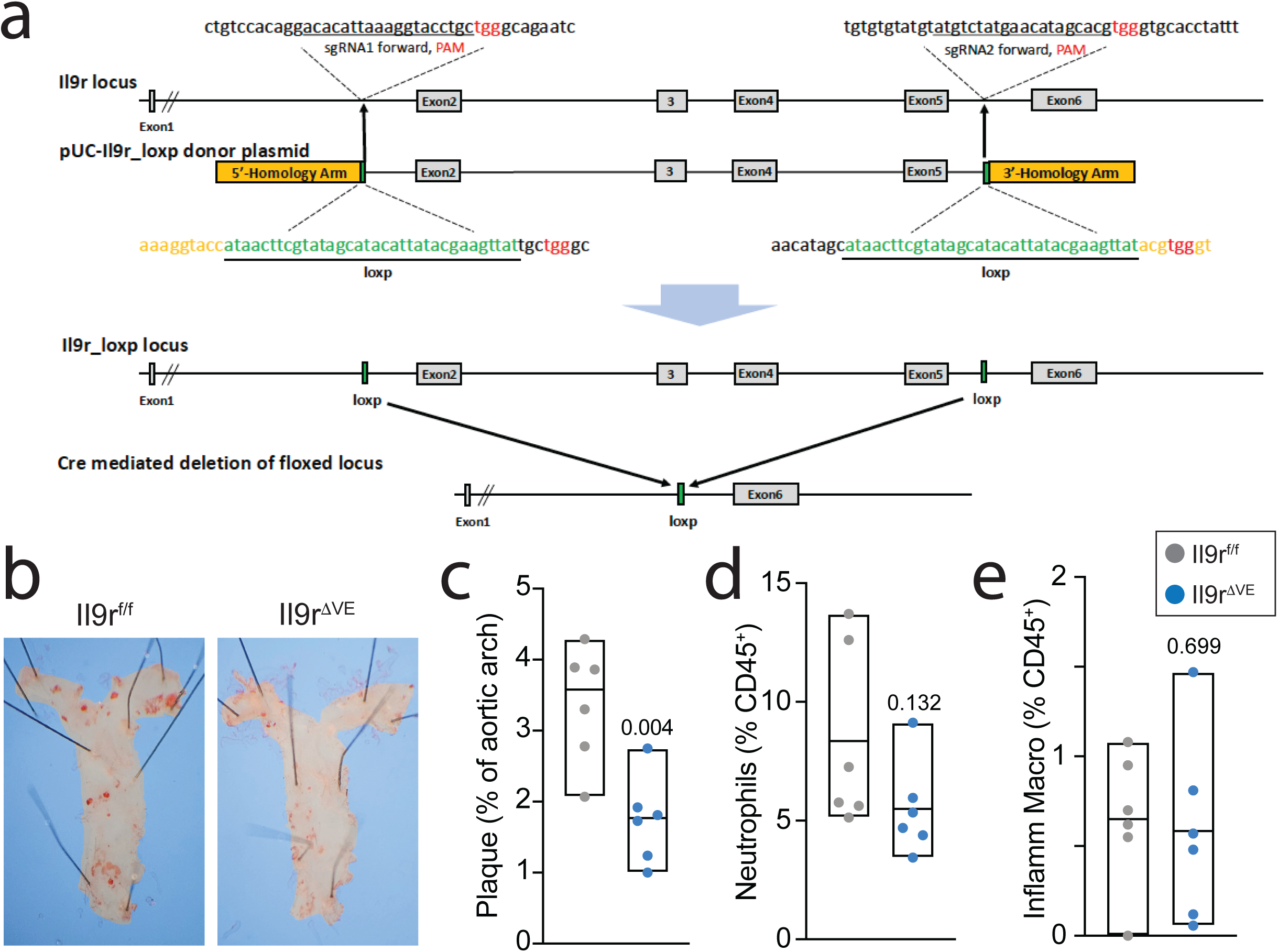
a. Generation of *Il9rf/f* animals using CRISPR/Cas9 editing. Schematic of mouse *il9r* locus, targeting construct of *il9rf/f*, and predicted knock-in allele. Two loxP sites were designed to surround the second and fifth exons, whose deletion resulted in a frame-shift mutation in exon6. The green boxes mark loxp sites, orange color represents the sequence of sgRNAs used and red indicate PAM sites. b,c. Representative images (b) and bar graph (c) show pooled oil red O (+) lipid-rich plaque as percentage of aortic arch in *ApoE-Il9^f/f^* (grey) and *ApoE-Il9^ΔVE^* mice (blue) exposed to three rounds of imiquimod (IMQ, Fig 4a**, S2j**) while being exposed to high-fat diet (HFD). e. Bar graphs show percent neutrophils (c) and Ly6C+ inflammatory macrophages in *ApoE-Il9^f/f^* (grey) and *ApoE-Il9^ΔVE^* mice (blue) exposed to IMQ-HFD. For all experiments: p-values, Mann-Whitney.

To delete IL-9 receptor in endothelial cells, we crossed *Il9r*^f/f^ mice to VE-Cre (*Cdh5*^Mlia^-Cre) mice (**Fig 6a**), which express Cre-recombinase in VE-Cadherin^+^ endothelial progenitor cells and show >95% recombination efficiency in mature endothelial cells (Alva et al., 2006). While VE-Cadherin is also expressed during hematopoiesis, *Cdh5*^Mlia^-Cre mediated deletion is inefficient in most leukocyte populations, allowing for preserved gene expression (Fahs et al., 2014). To induce psoriatic atherogenesis, we crossed *Il9r*^f/f^ x VE-Cre (henceforth termed *Il9r*^ΔVE^) mice to *ApoE*^-/-^ mice (henceforth termed *ApoE-Il9r*^ΔVE^). We then exposed *ApoE-Il9r*^ΔVE^ mice and *ApoE-Il9r*^f/f^ littermates to HFD for 6 weeks while treating with IMQ. Compared with *ApoE-Il9r*^f/f^ mice, *ApoE-Il9r*^ΔVE^ mice had significantly less lipid-rich aortic plaque (**Fig 6b,c**) and a trend towards reduced aortic infiltration of neutrophils (**Fig 6c**) but no difference in aortic infiltration of other leukocyte populations (**Fig 6d-e,S4d**) or in skin-infiltrating leukocyte populations (**Fig S4e**).

### IL-9 directly regulates arterial endothelial cells via STAT3

Our observation that deletion of IL9 receptor in endothelial cells prevented aortic plaque formation led us to hypothesize that IL-9 might directly target arterial endothelial cells to promote atherosclerotic disease. To test this hypothesis, we first set out to investigate IL-9 receptor expression in arterial endothelial cells. In 6 coronary arteries from 3 deceased subjects with a documented history of psoriasis, we found IL-9R+/VE-Cadherin+ cells surrounding lipid-rich plaque and at the arterial lumen (**Fig 7a-b**). The IL-9 receptor is heterodimeric molecule comprising the IL-9R and IL-2Rγ subunits (Schwartz et al., 2016). Both subunits were highly expressed and interacted in HAoEC, suggesting the presence of a functional IL-9 receptor (**Fig 7c-d**). IL-9 can signal through STAT1, STAT3, and STAT5 (Scharli et al., 2025): HAoEC expressed STAT1 and STAT3, (**Fig 7e**), and treatment with IL-9 induced phosphorylation of STAT3 (**Fig 7f,g**) but not STAT1 (**Fig 7h**). IL-9 treatment also induced expression of the STAT3 target gene *SOCS3* (**Fig S5a**)

**Fig 7.**
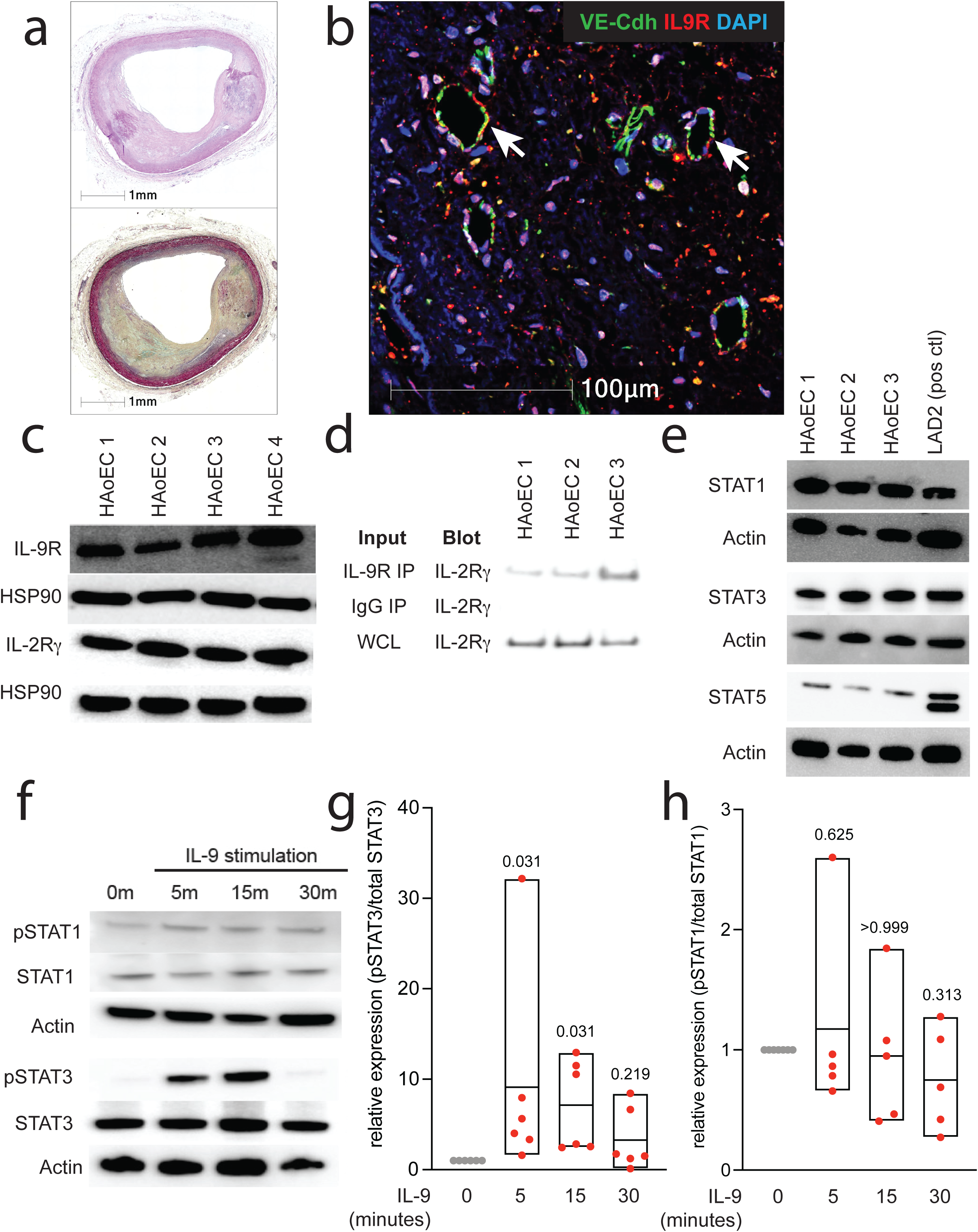
a-b. Representative images of coronary artery plaque from subject with psoriasis and atherosclerotic coronary artery disease show H&E and MOVAT staining (a) and immunofluorescence staining (b) for IL-9 receptor (red) and the endothelial marker VE-cadherin (green). c. Western blot shows expression of the IL-9 receptor subunits IL-9R and IL-2Rγ in primary aortic endothelial cells (HAoEC) from four cadaveric donors. d. Western blot shows expression of IL-2Rγ in whole cell lysates (WCL, positive control), lysates immunoprecipitated with IgG (IgG IP, negative control), and lysates immunoprecipitated with IL-9 receptor antibody (IL9R IP) in HAoEC from three cadaveric donors. e. Western blot shows expression of STAT1, STAT3, and STAT5 in HAoEC from three cadaveric donors and from the LAD2 mast cell line (positive control, known to express STAT1/3/5). f-h. Representative Western blot shows total and phosphorylated STAT1 and STAT3 expression in HAoEC treated with IL-9 and analyzed at various time points. Bar graphs show pooled quantification of phosphorylated/total STAT3 (g) and STAT1 (h) from HAoEC (n = 5 for STAT1, n = 6 for STAT3). For all experiments: p-values, Wilcoxon.

### IL-9 induces endothelial dysfunction through STAT3

Disruption of endothelial barrier integrity is a key step in the early pathogenesis of atherosclerotic cardiovascular disease(Pepin and Gupta, 2024), and STAT3 is thought to play a role in basal endothelial barrier function (Krempski et al., 2024). We, therefore, hypothesized that IL-9/STAT3 signaling might disrupt endothelial barrier integrity. To test this hypothesis, we treated HAoEC with IL-9 alone or in the presence of the STAT3 inhibitor C188-9. IL-9 disrupted endothelial monolayer morphology, significantly increasing the number and size of gaps between cells (**Fig 8a-b**). IL-9 also increased endothelial monolayer permeability as measured by electrical cell-substrate impedance sensing (ECIS) and by transwell monolayer permeability (Adil and Somanath, 2021) (**Fig 8c-d**). At the transcript level, IL-9 significantly induced *VCAM1* and *SELE*, and there was a trend towards *ICAM1* induction (**Fig S5b-d**). These genes are induced in pro-inflammatory states and are involved in leukocyte recruitment to lipid-rich plaque (Baumer et al., 2017; Pepin and Gupta, 2024). Together, these findings indicated that IL-9 directly induces endothelial dysfunction. C188-9 prevented IL-9-mediated disruption of endothelial integrity (**Fig 8a-d**), confirming that IL-9 promotes endothelial dysfunction through activation of STAT3.

**Fig. 8.**
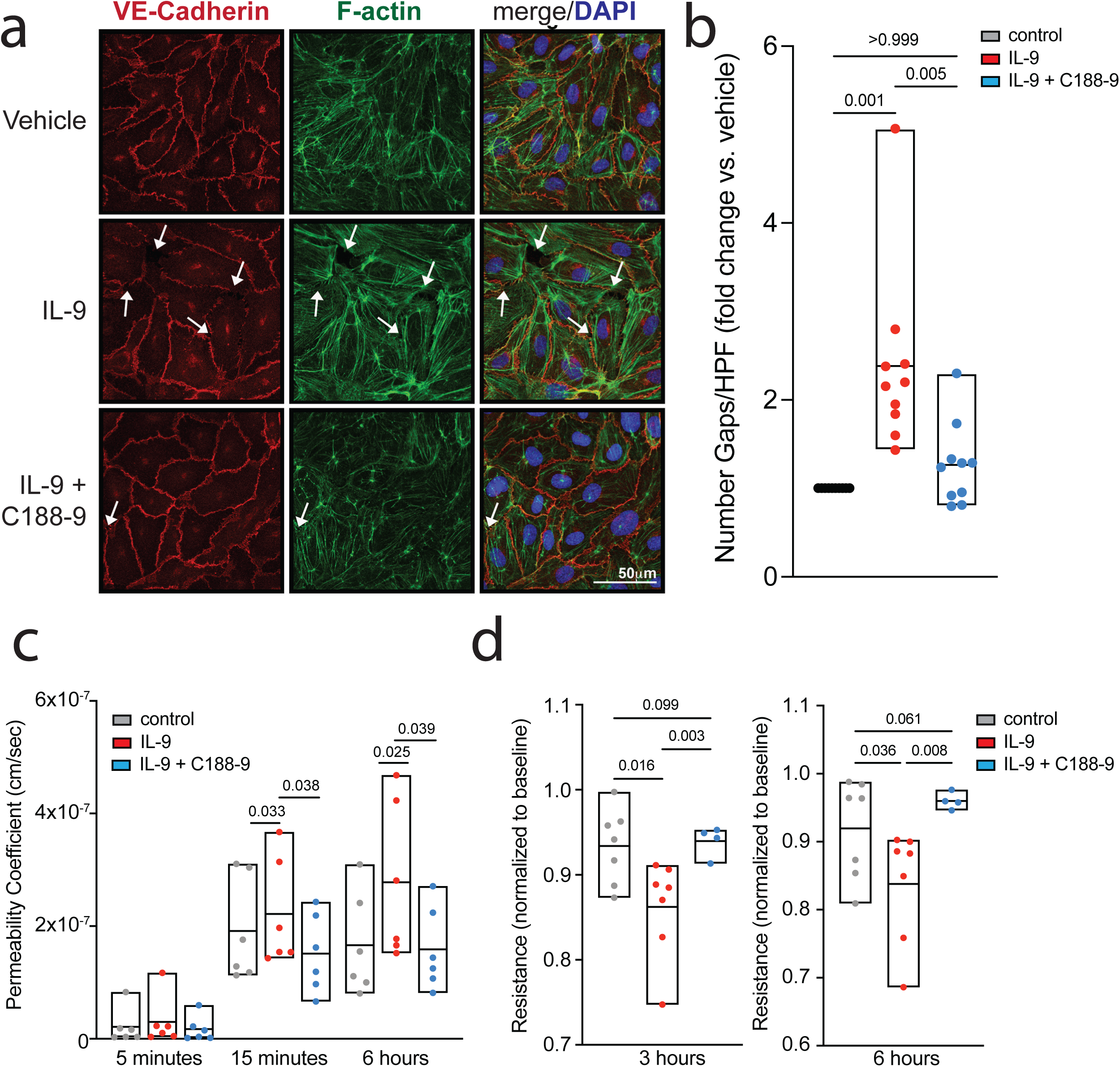
a. Representative image shows immunofluorescence staining of primary human aortic endothelial cells (HAoEC) in monolayer, treated with vehicle vs. IL-9 in the presence vs. absence of STAT3-inhibitor C188-9 and then stained for VE-Cadherin (red) and F-actin (green) to visualize monolayer integrity (Nuclei are displayed in blue, DAPI). b-d. Bar graph shows pooled results for number of gaps between cells (b), FITC-dextran permeability coefficient (c), or impedance measured by ECIS (e) for HAoEC treated with vehicle vs. IL-9 in the presence vs. absence of C188-9. For all experiments: p-values, ANOVA.

### IL-9 regulates adhesion and barrier-promoting genes in endothelial cells

STAT3 is a signal-dependent transcription factor that primarily exerts its effects via regulation of gene expression, inducing and repressing large cassettes of target genes (Schwartz et al., 2016). We, therefore, set out to determine the molecular targets of IL-9-STAT3 signaling at a global level by analyzing the transcriptomes of IL-9-treated HAoEC. IL-9 induced 278 genes (FC ≥ 2, adj p < 0.05, DESeq2), and repressed 681 genes (FC ≤ -2, adj p < 0.05, DESeq2) in HAoEC (**Fig 9a**). Because IL-9/STAT3 signaling disrupted endothelial barrier integrity, we hypothesized that IL-9 might reduce the expression of tight or adherens junction proteins. Although IL-9 modestly repressed *TJP2*, which links tight junction membrane proteins to cytoskeletal actin (Chistiakov et al., 2015), other genes involved in endothelial intercellular junction formation were upregulated by IL-9 (**Fig 9b**). These results indicated that IL-9 does not reduce barrier integrity by directly modulating the expression of junction-forming genes. To identify other potential mechanisms promoting endothelial dysfunction downstream of IL-9/STAT3, we analyzed enriched pathways in IL-9-induced and IL-9-repressed gene cassettes. Globally, IL-9-induced genes and IL-9-repressed gene cassettes were both enriched for proangiogenic, proinflammatory, and TGF-β signaling pathways (**Fig 9c,d**). Multiple pro-angiogenic pathways have been implicated in endothelial dysfunction downstream of STAT3, including Rho, VEGF, YAP-TAZ, and Smad signaling (Becerra et al., 2017; Shen et al., 2021; Wang et al., 2021; Wei et al., 2018). IL-9 modulated the expression of genes involved in all four pathways (**Fig 9e**). Together, these findings suggest that IL-9/STAT3 signaling modulates a broad cassette of genes that combine to promote endothelial barrier disruption and atherogenesis.

**Fig 9.**
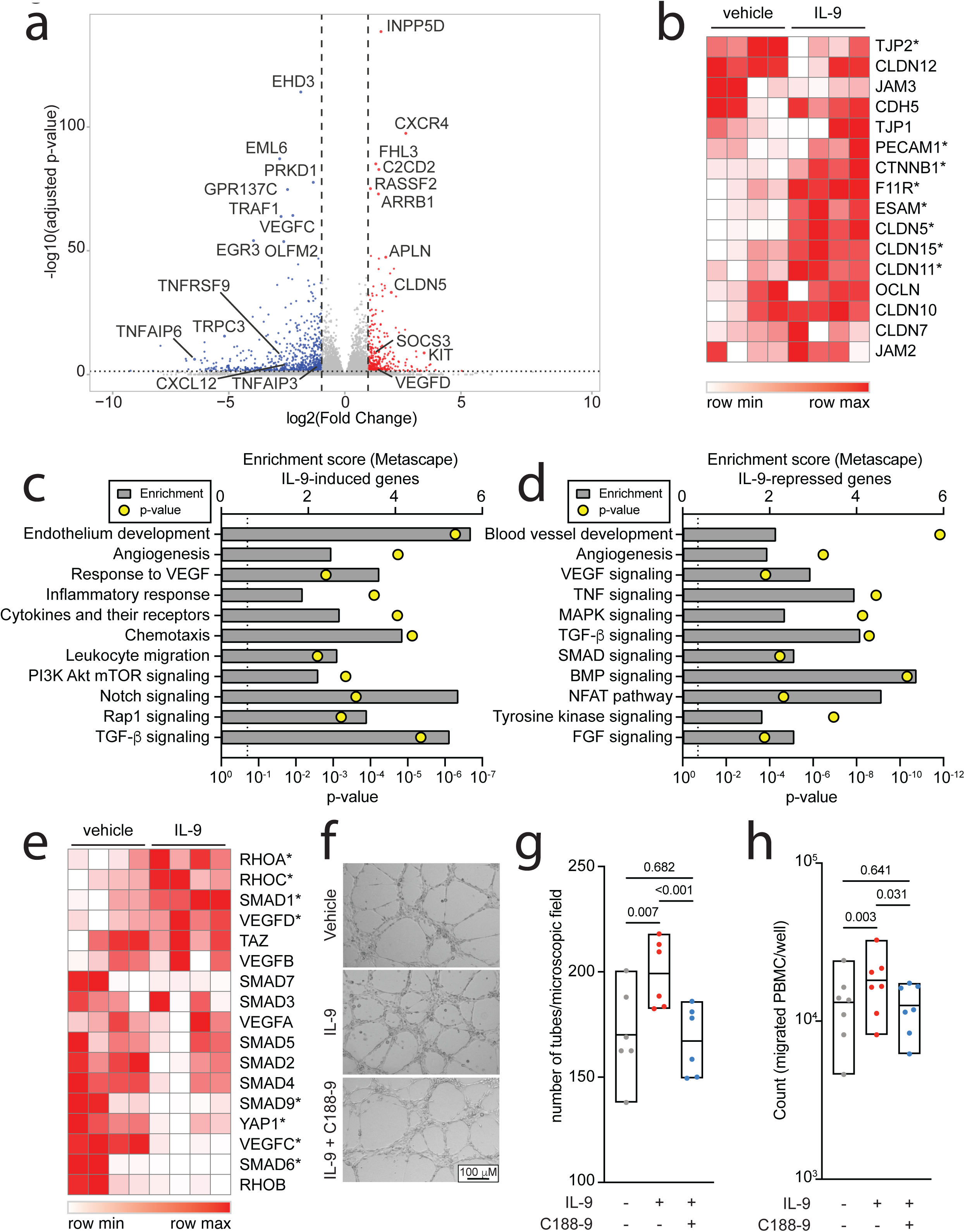
a. Volcano plot shows log_2_ normalized fold-change in gene expression vs. -log_10_ normalized Benjamini-Hochberg adjusted p-value (DESeq2) for primary human aortic endothelial cells (HAoEC, n = 4) treated with IL-9 vs. vehicle. Selected genes are marked. b. Heatmap shows expression (normalized across rows) of genes encoding tight and adherens junction proteins in endothelial cells treated with IL-9 vs. vehicle. * adjusted p-value <0.05, DESeq2. c,d. Bar graphs show enrichment scores (grey) and p-values (yellow) for selected enriched pathways (Metascape) in genes induced (c) or repressed (d) by IL-9 in HAoEC. b. Heatmap shows expression (normalized across rows) of genes encoding pro-angiogenic pathways associated with STAT3-mediated endothelial dysfunction, in endothelial cells treated with IL-9 vs. vehicle. * adjusted p-value <0.05, DESeq2.

### Endothelial IL-9R/STAT3 signaling enhances angiogenesis and leukocyte chemotaxis

Our transcriptomic analysis suggested that, in addition to endothelial barrier disruption, endothelial IL-9R/STAT3 signaling might promote leukocyte chemotaxis and angiogenesis. To test this hypothesis, we treated primary HAoEC with IL-9 in the presence vs. absence of C188-9, then measured *in vitro* endothelial 2D tube formation, a measure of angiogenesis (Ponce, 2009). IL-9-treated HAoEC formed more endothelial tubes (**Fig 9f,g**), confirming that IL-9 is pro-angiogenic in arterial endothelial cells. Supernatants from IL-9-treated HAoEC also promoted recruitment of healthy donor-derived human peripheral blood mononuclear cells (PBMCs) through a transwell (**Fig 9h**), suggesting that IL-9 stimulates release of leukocyte chemoattractant factors from HAoECs. Treatment with C188-9 reduced both PBMC migration and tube formation (**Fig 9f-h**). Together, these data suggest that IL-9R/STAT3 signaling induces a pro-chemotactic and pro-angiogenic program in HAoEC, thereby promoting endothelial dysfunction and inflammatory atherogenesis.

## Discussion

Inflammation is a major contributor to ASCVD, especially in patients with inflammatory and autoimmune diseases (Batty et al., 2022; Chistiakov et al., 2015; Doring et al., 2024; Eder et al., 2015; Guzman-Martinez et al., 2022; Harrington et al., 2017; Libby et al., 2002; Manning et al., 2025; Ridker et al., 2017; Wolf and Ley, 2019). However, major questions continue to surround the pathogenesis of inflammatory atherogenesis. These include the cytokines promoting inflammatory atherogenesis in patients with autoimmunity, as well as the sources and targets of these inflammatory cytokines. Here, we addressed several of these questions using transcriptomic, genetic, and functional approaches in murine models and primary human cells.

Psoriasis and other inflammatory diseases are characterized by expansion of Th9 cells, which can induce allergy, autoimmune inflammation, and antitumor immunity (Gerlach et al., 2014; Liu et al., 2014; Lu et al., 2018; Rauber et al., 2017; Schlapbach et al., 2014; Xue et al., 2019; Yao et al., 2013). Here, we find evidence that IL-9 produced by Th9 cells directly promotes inflammatory atherogenesis in subjects with psoriatic disease. IL-9 promotes atherogenesis in the context of multiple psoriatic disease models, including treatment with the TLR7 agonist imiquimod. Prolonged treatment with imiquimod can also induce a lupus-like phenotype, which includes elevated levels of autoantibodies and vasculopathy (Liu et al., 2018; Yokogawa et al., 2014). This suggests that IL-9 may be key driver of inflammatory and autoimmunity-related CVD beyond psoriatic disease. Indeed, IL-9 and Th9 cells have previously been implicated in atherosclerosis and myocardial infarction (Li et al., 2017; Zhang et al., 2015), and we detect *SPI1*^+^ cells expressing a Th9 transcriptional program in the atherosclerotic plaque of non-psoriatic subjects. Our findings therefore suggest that psoriasis and other Th9-high diseases increase this pro-atherogenic state, leading to earlier onset of CVD. Our findings could also help explain why asthma and atopic dermatitis are associated with increased CVD risk (Guo et al., 2022; Wan et al., 2023), although the “type 2” allergic cytokines IL-4, IL-5, and IL-13 do not promote atherogenesis and may even be protective (Binder et al., 2004; Cardilo-Reis et al., 2012; King et al., 2007). Future studies will be needed to specifically address the role of Th9 cells and IL-9 in asthma- and allergy-associated atherogenesis and CVD risk.

Our results also answer important questions about Th9 lineage identity in the context of various disease states. Previous studies have suggested considerable heterogeneity of Th9 cells across different tissues, disease, states, and immunological contexts (Kaplan, 2017; Kaplan et al., 2015; Licona-Limon et al., 2013; Schwartz et al., 2019; Seumois et al., 2020; Ulrich et al., 2022). We previously identified an “allergic Th9” transcriptional program that was conserved across human subjects and murine allergy models (Son et al., 2023). We now find enrichment of the “allergic Th9” cassette in circulating and tissue-resident cells from subjects with autoimmune disease, definitively establishing that Th9 cells have a conserved lineage identity that spans affected tissues and diseases. Analysis of “psoriatic Th9” cells reveal upregulation of markers promoting migration to the heart and into atherosclerotic vessels, which could help to explain the presence of plaque-infiltrating Th9 cells detected in scRNAseq and immunohistochemical analysis of human carotid and coronary samples. Previous work has suggested that Th9 cells in patients with psoriatic disease initially differentiate in the skin or small intestine, then enter the circulation and migrate to distal sites to promote systemic inflammation (Ciccia et al., 2016; Schlapbach et al., 2014). Conversely, atherosclerosis is characterized by CD4+ T cell responses to viral epitopes and to apolipoprotein B (ApoB), the main lipoprotein in LDL (Chowdhury et al., 2022; Saigusa et al., 2022). Although the mechanisms promoting generation of ApoB-reactive T cells are poorly understood, psoriasis and atherosclerosis are both associated with inflammation of visceral adipose tissue, suggesting that plaque-resident Th9 cells may differentiate in the visceral fat (Ohman et al., 2008; Schwartz et al., 2022). A comprehensive understanding of the origin and migration of plaque-infiltrating Th9 cells will require future investigation and methodologies, including fate-reporter and dynamic tracking mouse models.

Prior studies have suggested that Th9 effector function is transient without sustained activation from external factors. Our findings indicate that multiple pro-atherogenic factors regulate Th9 differentiation, which could be a possible mechanism for sustained or restimulated IL-9 production within the atherosclerotic plaque. Previous studies have demonstrated that cytokines and mediators found in atherosclerotic plaque can profoundly influence Th9 lineage specification: these include IL-1β, hypoxia, nitric oxide, and intracellular fatty acids (Canaria et al., 2022; Nakajima et al., 2024; Niedbala et al., 2014; Roy et al., 2021; Xue et al., 2019). We now find that VEGF-A and oxidized LDL have significant yet disparate effects on Th9 differentiation, adding to the complex network of Th9-regulated factors and cytokines. Future studies will be needed to determine the precise mechanisms by which VEGF-A and other proatherogenic factors combine to modulate Th9 differentiation and expansion during inflammatory atherogenesis.

In addition to resolving some of the questions surrounding Th9 identify and function, our studies also uncover a direct role for IL-9/IL-9R/STAT3 signaling in endothelial cells. This finding is particularly interesting because IL-9 signals through a heterodimer of IL-9R and IL-2RG, or the common gamma chain (γc) (Pajulas et al., 2023). Most γc cytokines are thought to primarily exert their effects by regulating the actions of hematopoietic cells, and most non-hematopoietic cells do not express IL-9R or IL-2RG under homeostatic conditions, although allergic inflammation can induce IL-9R expression (Pajulas et al., 2023). IL-9R expression has been seen in inflamed human coronary arteries (Ohtsuki et al., 2023), murine aortic endothelial cells (Zhang et al., 2015), and human atherosclerotic plaque-infiltrating immune cells (Gregersen et al., 2013). This widespread IL-9R expression has made it difficult to determine the role of endothelial IL-9R/STAT3 signaling in promoting inflammatory atherogenesis. Our results directly address this question demonstrating that IL-9 directly targets human arterial endothelial cells to induce dysfunction through diverse mechanisms. In addition to endothelial cells, IL-9R signaling may also modulate leukocyte migration and function to promote atherogenesis; one limitation of endothelial-directed Cre recombinases including VE-Cre is that VE-cadherin is expressed during hematopoiesis, leading to deletion of target genes like IL-9R in some leukocyte subsets. IL-9/IL-9R signaling is important for pulmonary macrophage recruitment and expansion in allergic asthma (Fu et al., 2022), suggesting that it could also promote macrophage recruitment in inflammatory atherogenesis – a key pathogenic mechanism (Libby et al., 2002). Indeed, chemotaxis genes were enriched within the cassette of genes induced by IL-9 in endothelial cells; hence, IL-9 may directly and indirectly promote leukocyte recruitment in addition to inducing endothelial dysfunction.

In addition to transducing signals downstream of IL-9/IL-9R, STAT3 is also important for signaling downstream of pro-atherogenic cytokines like IL-6 and VEGF-A, as well as protective cytokines like IL-10 (Short et al., 2022; Wang et al., 2021; Wei et al., 2018). Accordingly, STAT3 has complex and context-dependent roles in endothelial biology. STAT3-deficient endothelial cells are unable to suppress interferon gamma, which is protective in the context of toxic shock (Kano et al., 2003). However, STAT3 induces vascular permeability, angiogenesis, and extracellular matrix remodeling under homeostatic conditions (Dmitrieva et al., 2020; Hox et al., 2016). In addition to promoting homeostatic endothelial barrier function, our work suggests that STAT3 also promotes endothelial barrier dysfunction downstream of IL-9 signals through STAT3 to promote endothelial cell dysfunction through induction of a multifunctional gene cassette.

Finally, our findings open novel and promising avenues for immunomodulation to prevent CVD by blocking IL-9 or its receptors. Most other cytokines linked to inflammatory atherogenesis have critical roles in host-pathogen defense. Hence, therapeutic blockade increases the risk of infection – this can significantly counteract the benefits of immunomodulation for CVD prevention (Mitsis et al., 2024). By contrast, IL-9 is primarily involved in defense against helminths (Licona-Limon et al., 2017), which are not prevalent in developed countries. Early phase trials using IL-9-blocking antibodies for asthma have not shown any warning signal for infectious disease risk (Oh et al., 2013; Parker et al., 2011), and type 2 cytokine blockers are generally considered safer than other biologics (Mansilla-Polo and Morgado-Carrasco, 2024). In patients with concurrent autoimmunity and allergy and eosinophilia, type 2 blocking agents can even be added to other immunomodulators (Bettiol et al., 2022; Gisondi et al., 2023; Sanchez-Alamo et al., 2023). Although further studies are needed, these results suggest that IL-9 blockade could represent a safe and effective form of immunomodulation for CVD prevention.

In summary, IL-9^+^ Th9 cells are expanded in patients with the autoimmune disease psoriasis and are significantly associated with non-radiographic coronary artery disease. Psoriatic Th9 cells express a conserved Th9 transcriptional program that poises them to home to the heart and infiltrate blood vessels, where they are modulated by pro-atherogenic factors. Within the vessel, Th9 cells release IL-9, which directly targets endothelial cells via IL-9R/STAT3 signaling to induce endothelial dysfunction and worsen inflammatory atherogenesis. Together, these results uncover a novel mechanism of inflammatory atherogenesis that presents a promising therapeutic target in patients with autoimmunity.

## Methods

### Human Samples and Data

Whole blood, coronary artery CT scores, and demographic parameters were obtained from de-identified patients with psoriasis who were evaluated at the NIH Clinical Center under the Institutional Review Board (IRB) approved protocol 13H-0065 (Harrington et al., 2017). Samples were obtained and analyzed from untreated patients as well as patients taking immunomodulatory medications including biologic disease-modifying antirheumatic drugs. Full details of the protocol are reported elsewhere. De-identified buffy coat samples and whole blood samples from non-psoriatic volunteers were obtained from the NIH blood bank or through a separate IRB-approved protocol at the NIH clinical center (NCT019344660). All human atherosclerotic coronary artery specimens from psoriasis patients were obtained from the CVPath autopsy registry; this registry study was approved by the Institutional Review Board (IRB) at CVPath Institute.

### Mouse experiments

Experiments were conducted on 8–12-week-old mice that were age- and sex-matched within experiments. All experiments were conducted in male and female mice and were performed in an Association for Assessment and Accreditation of Laboratory Animal Care-accredited animal facility, approved by the National Institute of Allergy and Infectious Diseases (NIAID) Institutional Animal Care and Use Committee (IACUC), or by the University of Pittsburgh IACUC. Experiments were performed in accordance with NIH guidelines for the use and care of live animals, under protocol LAD12E (NIAID) or protocol 22070649 (University of Pittsburgh). Mice were maintained in a specific-pathogen-free facility on a 12-h light–dark cycle, in open cages. Food and water were continuously available, except for (in high fat diet experiments) the 24h prior to euthanasia. Mice were checked periodically to ensure normal health and more often if adverse effects were anticipated (i.e. in psoriasis models). VE-Cre (*Cdh5-Mlia*), *Card14^ΔE138^*, and *ApoE^-/-^* mice were purchased from The Jackson Laboratory. *Il9r^-/-^* mice were generously provided by M Kaplan and J-C Renauld. INFER mice were generously provided by R. Flavell and P. Licona Limon. All mice were on a C57BL/6 background.

### Generation of *Il9r*^f/f^ mice

*Il9r^f/f^* mice were generated by CRISPR/Cas9-method at the Mouse Genetics and Gene Modification Section, Comparative Medicine Branch, NIH. Gene targeting was aimed to delete from exon2 to 5 of il9r gene and the locations of flox insertions were carefully chosen not to disturb proper splicing of il9r (Fig. x). A 3.5kb DNA cassette including 800bp homology arms at both sides was synthesized and inserted into pUC vector by Azenta Life Sciences (Burlington, MA). Guide RNAs were selected using CRISPOR design tools (http://crispor.tefor.net/), and synthetic gRNA was manufactured by Synthego (Redwood City, CA). Cas9 nuclease was purchased from IDTdna (Coralville, IA). For microinjection, the C57BL/6Tac females were injected with 7.5IU of pregnant mare serum gonadotropin (Prospec, Israel) followed by 5IU of human chorionic gonadotropin (Sigma, St. Louis, MO) after 47 hours, then mated with C57BL/6Tac males. The following morning, fertilized one-cell-stage embryos were collected, washed 6 times with EmbryoMax M2 media (Millipore Sigma, Burlington, MA), and then cultured in EmbryoMax KSOM media (Millipore Sigma, Burlington, MA) at 6% CO_2,_ 37°C for 3 hours or until the microinjection. The embryos were transferred to 50mm glass bottom dish (Mattek, Ashland, MA) in EmbryoMax M2 drop, overlaid with embryo-tested mineral oil (Sigma, St Louis, MO), and microinjected using an DMi8 inverted microscope (Leica, Wetzlar, Germany) equipped with a set of TrasferMan4 micromanipulator and FemtoJet4i (Eppendorf, Hamburg, Germany). Microinjection mixture was prepared fresh as following: 10ng/µl of HiFi Cas9 protein, 5ng/µl of sgRNA, 5ng/µl of plasmid in microinjection buffer (10mM Tris-HCL pH7.5 with 0.25mM EDTA), backfilled to microinjection needles for pronuclear injection. Injected embryos were implanted into the oviducts of pseudo-pregnant surrogate CD-1 females (Charles River Labs, Wilmington, MA) after 1 hour culture in EmbryoMax KSOM at 6% CO_2,_ 37°C. Offspring born to the foster mothers were genotyped by PCR. The primer sequences for genotyping were as follows: il9r for, CACAAGTCCGAGTTGAGGTC; il9r 5flox rev, AATGTATGCTATACGAAGTTATGGTACC; il9r 3flox for, AGCATACATTATACGAAGTTATACGTG; il9r rev, GGATGCTGGCTGACCATT.

### Murine psoriasis models

Three murine models of psoriasis were used. For the Imiquimod (IMQ)-ApoE model (Fig 4a), 9-11 week old *ApoE*^-/-^ mice were shaved on their dorsal caudal/posterior bodies and then treated with daily applications of 60mg Imiquimod 5% cream (IMQ, Glenmark pharmacy) or Vanicream (Pharmaceutical Specialtis, Inc) for three 5-day cycles, with 13-day rest periods between applications to allow skin recovery (6 weeks in total). For the IL-23-ApoE model, *ApoE^-/-^* mice were shaved on their flanks and then treated with intradermal injections of IL-23 (carrier free) recombinant protein (ebioscience, 34-8231-82) 500ng in 20ul PBS per day for two 5-day cycles, with 59-day rest periods between injection (10 weeks in total). For the Card14-ApoE model, Card14^ΔE138^ mice were crossed to *ApoE ^-/-^* mice and developed spontaneous psoriasis.

For induction of atherogenesis, mice were exposed to a high-fat diet (HFD, 42% fat) Envigo, TD.88137. For the IMQ and IL-23 (induced) psoriasis models, mice were exposed to HFD starting on d1 of psoriasis induction. HFD was continued for 6 weeks (IMQ) or 10 weeks (IL-23), or throughout psoriasis induction. For the Card14 model (spontaneous psoriasis), mice were exposed to HFD starting at 8-12 weeks of age and continued for 7 weeks.

In some cases, mice were injected with mouse IL-9 blocking antibody (BioXCell, BE0181) or mouse IgG2a isotype control (BioXCell, BE0085) at 100μg in 200ul two times a week. for the IMQ and IL-23 (induced) psoriasis models, IL-9-blocking antibodies were used starting on d1 on psoriasis induction and continued until mice were euthanized. For the Card14 model (spontaneous psoriasis), mice were exposed to IL-9-blocking antibodies starting on d1 of HFD feeding, and treatment was continued until mice were euthanized.

### Skin digestion

Mouse dorsal caudal/posterior skin (∼3cm X 4cm) was cut out and digested as previously described (Sakamoto et al., 2022). Briefly, the subcutaneous layers from all the excised skin samples were removed, and the skins were placed in PBS. After removing the excess PBS, each skin was cut into tiny pieces with scissors. The skin was placed on ice and further minced with the addition of 1 ml skin digestion buffer (10μl of 25mg/ml (0.65 Wunsch units/mL) Liberase TL in PBS). The minced skin and the digestion buffer were transferred to a 6 well plate containing 4ml digestion buffer, then incubated at 37°C for 2 hours, with the addition of 1ml Trypsin for the last 10 minutes. The reaction was stopped by adding 4-5ml 5% FACS buffer. Samples were mixed and passed through 100μm filters, then centrifuged (400g x 8 min, 4°C). Cells were then resuspended in FACS buffer and passed through 40μm filters, then stimulated and/or stained as below.

### Aorta digestion

Aorta samples were cut in 1mm rings and were collected in a 15ml Falcon tube containing 200μl aorta digestion buffer containing 10μl of 25mg/ml (0.65 Wunsch units/mL) Liberase TM in PBS. The tubes were incubated in a 37°C water-bath for 40 minutes, resuspending them every 15 minutes. The reaction was stopped by the addition of 10ml PBS with 2% FBS. The contents of the 15ml tube were passed through 40μm filters and then centrifuged at 1400 rpm for 5 minutes. After decanting the supernatant cells were resuspended and processed for staining. After digestion, cells were stimulated and/or stained as below.

### Spleen digestion

Mouse spleens were collected in 1ml skin digestion buffer and kept on ice until all the samples were harvested. Each spleen was mashed over a 40μm filter with 20ml spleen digestion buffer consisting of 10% FBS and 1μg/ml DNase I and collected in a 50ml tube. The tubes were centrifuged at 1400 rpm, for 5 minutes, and supernatants were decanted. Cell pellets were resuspended in 2ml ACK lysis buffer and incubated for 3 minutes to deplete RBCs. The reaction was stopped by the addition of 10ml of spleen digestion buffer.

### Aortic plaque measurements (mice)

After sacrifice to assess murine atherosclerosis development in the aorta, the vasculature was perfused with PBS, and the entire aorta was cleaned from adventitial fat under a dissection microscope. Subsequently, the thoracic part of the aorta was utilized for *en face* atherosclerotic lesion quantification while the abdominal part of the aortas were used for flow cytometry. For *en face* atherosclerotic lesion analysis, the thoracic part of the aortas were cut open longitudinally, transferred to a wax-coated tray, and pinned using micro pins (26002-10, Fine Science Tools) under a dissection scope. Afterwards, the aortas were fixed using 4% paraformaldehyde solution for 10 minutes. To visualize lipid-rich atherosclerotic plaques, Oil Red O staining was performed as described previously (Baumer et al., 2017). Briefly, after fixation aortas are washed with water and incubated with isopropanol (60%) for 5min. Subsequently, the Oil Red O solution was added and incubated for 15 min under slow rotation. Excessive Oil Red O solution was washed away in three 5 min washing steps with 60% isopropanol. For imaging the aortas were submerged in water. Photos of the sections taken at 10x magnification were analyzed for Oil Red O positive area (ImageJ) and expressed as the percentage of total aortic area.

### In vitro human and murine Th9 differentiation

Naïve human and murine T cells were enriched to >90% purity using magnetic separation, as described previously(Son et al., 2023). Human naïve T cells were plated at a density of 1e6 per mL in 1mL in a 48-well plate, on plate bound αCD3 (OKT3 1μg/mL). Cells were cultured in complete RPMI-1640 with glutamine (10% FBS, 100 IU/ml, penicillin, 0.1 mg/ml streptomycin) in the presence of cytokines and antibodies to promote the differentiation of Th9 cells (1 μg/ml αCD28, 2.5 μg/ml αIFN-γ, 5 ng/ml hTGF-β, 10 ng/ml hIL-2, 30 ng/ml hIL-4, 10 ng/ml hIL-1β). Murine naïve T cells were cultured in complete IMDM (10% FBS, 100 IU/ml, penicillin, 0.1 mg/ml streptomycin, 55 μM beta-mercaptoethanol) at a density of 1e6 cells per ml in 1mL in a 24-well plate. Cells were activated with plate-bound αCD3L (10μg/mL) and αCD28 (10 .g ml−1) for 3d with polarizing cytokines and antibodies to promote Th9 differentiation (10μg/mL αIFN-γ, 20 ng/ml mIL-4, 10 ng/mL hIL-2, 5 ng/ml hTGF-β). For some experiments, cells were cultured in the presence of vehicle vs. VEGF-A (various concentrations, Peprotech) or oxidized LDL (various concentrations, Lee Biosolutions 360-31). For oxidized LDL experiments FBS charcoal stripped (Gibco) was used. After 3d (mouse) or 5d (human) cells were stimulated and analyzed as below.

### Restimulation

All cells were stimulated in 96-well plates. Human PBMCs were isolated by density centrifugation (Ficoll-Paque) and cryopreserved. For restimulation assays, cells were thawed and washed with complete RPMI-1640 with DNAse as previously described, then restimulated at 37°C with 10 ng/mL phorbol 12-myristate13-acetate (PMA), 1µg/mL ionomycin, 20 ng/mL hIL-2, and 60 ng/mL hIL-4. After 2 hours, 10 μg/mL brefeldin was added for 4h (total 6h stimulation).

Murine *in vitro* differentiated Th9 cells were stimulated with 100 ng/mL PMA, 1 µg/mL ionomycin; 20 ng/mL hIL-2; 40 ng/mL mIL-4. Human *in vitro* differentiated Th9 cells were stimulated with 20 ng/mL PMA; 1 µg/mL ionomycin; 20 ng/mL hIL-2; 60 ng/mL hIL-4. In both, 10 μg/mL brefeldin was added after 2h, and the incubation was continued for another 4h (total 6h stimulation).

For stimulation of murine *in vivo* differentiated Th9 cells (skin, spleen, aorta): organs were digested as above, then cells were resuspended in cIMDM for 5 hours with PMA (100ng/mL), ionomycin (1 µg/mL), hIL-2 (20 ng/mL), and mIL-4 (40 ng/mL).

### Flow cytometry

Human PBMC were stained for viability (Live-Dead BLUE, Invitrogen), then fixed and permeabilized (BD Cytofix/Cytoperm). Cells were then stained with: CD3 APC-Cy7, CD4 BUV395, CD8 v500, CD45RO PE-Texas Red, IL-2 PerCp/Cy5.5, IL-4 BUV605, IL-13 BV711, IL-9 PE, IL-5 v450, IFN-γ AF700, TNF-α PE-Cy7, IL-17A FITC, IL-21 AF647, IL-10 e655. Data were collected on a LSRFortessa (BD) and analyzed with FlowJo (Treestar) and Prism (GraphPad).

In vitro differentiated murine Th9 cells were stained for viability (Zombie AQUA, BioLegend), then fixed and permeabilized (BD Cytofix/Cytoperm). Cells were then stained with CD4 PerCp/Cy5.5, IL-9 APC, IFN-γ PE, Foxp3 v450, IL-4 AF488. Data were collected on a LSRFortessa (BD) and analyzed with FlowJo (Treestar) and Prism (GraphPad).

For analysis of tissue-infiltrating murine cells (*ex vivo*), suspensions of digested organs were transferred to 96-well plates and centrifuged (400g x 3min, 4°C), then washed (1X PBS) and stained for viability (Zombie AQUA, Biolegend) in the presence of Fc block. After 10 minutes, the following antibodies were added for 30 minutes at 4°C: F4/80 SB600, CD11b FITC, SiglecF PE, Ly6G PerCP/Cy5.5, Ly6C AF647. Cells were fixed and permeabilized (BD Cytofix/Cytoperm), followed by intracellular staining for 3 hours: CD3 BV421, TCRβ APC-Cy7, CD45.2 PE-Cy7, CD4 PE/Dazzle 594, CD8a AF700. In some cases, cells were stimulated for analysis of *in vivo* differentiated T cells and stimulated as above, then stained for viability (Zombie UV, Biolegend) in the presence of Fc block. After 10 minutes, the following antibodies were added for 30 minutes at 4°C: CD11b-FITC, Ly6C-Alexa Fluor 700, F4/80-SB 600, followed by intracellular staining for 2 hours: CD45APC-Cy7, CD3-Brilliant Violet 421, TCRβ-SB 645, CD4-PerCP-Cy5.5, IL9-PE, IL13-PE-Cy7, IL17-APC, IFNγ-Pacific Blue. Data were collected on a Cytek Aurora and analyzed with FlowJo (Treestar) and Prism (GraphPad). For analysis of Th9 cells from INFER mice, suspensions of digested organs were transferred to 96-well plates and stained with DAPI and cell surface antibodies in the presence of Fc block (20 μg/mL), heat-inactivated rat serum (1:50 dilution, ThermoFisher 10170C), and heat-inactivated hamster serum (Abcam ab7483). Cells were stained with: CD4 PerCpCy5.5, CD8 PE, TCR-β APC-Cy7, CD45.2 PE-Cy7, and CD44 AF700. Cells were sorted for CD45.2+, TCR-β+, CD4+, CD8-, CD44+ cells, then for GFP+ (Th9) and GFP-(non-Th9) cells.

### Immunofluorescent staining of human coronary arteries

Following perfusion fixation with 10% neutral buffered formalin, Human atherosclerotic coronary arteries were excised from the heart and subsequently decalcified using EDTA. The major epicardial arteries were segmented at 3 mm intervals, dehydrated, and embedded in paraffin blocks. Tissue sections were cut on a rotary microtome at 4-6 µm and stained with Hematoxylin and Eosin (H&E) or Movat Pentachrome for histological analysis.

For immunofluorescent staining, serial sections were deparaffinized in xylenes, rehydrated to distilled water, and subjected to heat-induced antigen retrieval prior to incubation with primary antibodies against CD3 (0.2μg/mL, 790-4341, Roche Diagnostics, Indianapolis, IN, USA) or CD4 (1.25μg/mL, 790-4423, Roche Diagnostics), co-labeled with PU.1 (1:400, 2258, Cell Signaling Technology, Danvers, MA, USA) overnight at 4 °C. Subsequently, sections were incubated with Alexa Fluor 488-conjugated anti-mouse IgG (1:150, A21202, Thermo Fisher Scientific, Waltham, MA, USA) and Alexa Fluor 555-conjugated anti-rabbit IgG (1:150, A31572, Thermo Fisher Scientific) secondary antibodies, respectively, for 2 hours at room temperature, followed by DAPI (1:500, D3571, Thermo Fisher Scientific).

For double staining of VE-cadherin and IL-9R, sections were incubated with antibodies against VE-cadherin (1:100, AF1002, R&D Systems, Minneapolis, MN, USA) and IL-9R (1:400, 310408, BioLegend, San Diego, CA, USA), followed by Alexa Fluor 488-conjugated anti-goat IgG (1:150, A11055, Thermo Fisher Scientific) and Alexa Fluor 555-conjugated anti-mouse IgG (1:150, A31570, Thermo Fisher Scientific), respectively, and DAPI (1:500, D3571, Thermo Fisher Scientific). Histological images were scanned at 10X magnification (AxioScan Z1, Zeiss) and immunofluorescent images were acquired using a laser-scanning confocal microscope (LSM800, Carl Zeiss Microscopy, Jena, Germany) with optical sectioning along the z-axis at 20X magnification.

### Endothelial cell culture, microscopy, immunofluorescence anaysis, ECIS, FITC-dextran

Primary human aortic endothelial cells (HAoEC) from ≥ 9 deceased individuals were purchased from PromoCell, Germany. HAoEC were kept in culture and passaged as described previously for up to 9 passages (Baumer et al., 2020; Baumer et al., 2017). Various assays were performed when HAoEC monolayers reached full confluency to assess endothelial function. All experiments were performed as >4 biological replicates.

Immunofluorescence analysis was utilized to visualize endothelial monolayers and quantify intercellular gap formation as described previously (Baumer et al., 2017). HAoECs were incubated with treatments as indicated upon reaching full confluency. After 6 hours of treatment, HAoEC monolayers were fixed with 2% paraformaldehyde in PBS with Ca/Mg for 5 min at room temperature, washed three times with PBS with Ca/Mg, and permeabilized for 5min at room temperature using 0.1% TritonX-100 in PBS with Ca/Mg. Subsequently, cells were washed again three times with PBS with Ca/Mg, blocked for 1 hour at room temperature using 5% normal goat serum in 2% BSA in PBS with Ca/Mg. Afterwards, monolayers were incubated with anti-VE-cadherin (1:100, clone F-8, Santa Cruz, USA) in 2% BSA in PBS with Ca/Mg overnight at 4°C. The next day monolayers were washed with PBS with Ca/Mg and incubated with Phalloiding-AF488 to label F-actin and DAPI to label nuclei. Images for gap quantification of 2-3 microscopic fields were taken with the EchoRevolve microscope. Intercellular gap presence was quantified using ImageJ. High-resolution images were taken at the NHLBI Light Microscopy Core using the 780 Zeiss Microscope. Endothelial barrier integrity was determined on confluent HAoEC monolayers using Electric Cell-substrate Impedance Sensing (ECIS) technology as described previously (Baumer et al., 2017). Additionally, we measured endothelial barrier function by measuring 70kDa FITC-dextran flux through HAoEC monolayers cultured on 0.4μm transwell filters as described previously (Baumer et al., 2010).

To analyze a potential impact of IL-9 on endothelial tube formation, the MatrigelTM tube formation assay was used as described previously (Baumer et al., 2010). Briefly, HAoECs were treated as indicated overnight. Afterward, supernatants were collected and treated HAoECs removed from culture surfaces for subsequent tube formation assay. The assay was performed two-fold: a) utilizing the collected supernatants on control HAoECs, and b) utilizing the treated HAoECs with normal growth media allowing tube formation for 4 hours. Images were taken with the EchoRevolve microscope, and the number of tubes per microscopic field was quantified using ImageJ.

To determine whether HAoECs treated with IL-9 might release factors to support immune cell chemotaxis, HAoECs were treated as indicated for 6 hours, and the supernatants were collected. PBMCs (2×10^6^ per condition) from healthy blood bank donors were seeded in the top of a 5μm transwell filter in a 24-well plate. The lower chamber was filled with 300μl of the treated HAoECs supernatants. PBMCs were allowed to transmigrate to the lower chamber for 4 hours. Afterward, the number of migrated cells was counted in the bottom chamber.

### mRNA extraction, RT-qPCR

Cells were suspended in Trizol LS (Invitrogen). Total RNA was extracted according to the manufacturer’s instructions and treated with DNAse to remove genomic DNA, then 500 ng RNA was reverse transcribed using a complementary DNA synthesis kit (Applied Biosystems) and quantitative PCR (qPCR) was performed in triplicate using TaqMan master mix (Applied Biosystems).

For gene expression in primary aortic endothelial cells, we used 8 biological replicates from 7 individual donors. The following primers were used to detect mRNA expression in hAoEC: human TBP (qHsaCIP0036255), VEGFA (qHsaCEP0050716), VCAM1 (qHsaCIP0030799), ICAM1 (qHsaCEP0024986), SELE (qHsaCIP0028854), SOCS3 (qHsaCEP0024741). Expression of transcripts was normalized to TBP.

### mRNA sequencing library preparation

RNA was extracted as above. Libraries for mRNA-sequencing were prepared using the NEBNext Ultra II Directional RNA-seq Library kit, as per the manufacturer’s instructions. Samples were sequenced on a NovaSeq 6000, paired end (PE), for 50 cycles and processed with bcl2fastq v.2.20.0.422

### mRNA sequencing analysis

Reads of 50 bases were processed using the University of Pittsburgh nextflow-core pipeline (Ewels et al., 2020) (https://nf-co.re/rnaseq/2.0). Briefly, adaptor sequences were trimmed with Trim Galore and mapped to the mouse transcriptome and genome mm10 or human transcriptome and genome GRCh38 (hg38) using STAR. Gene expression values (fragments per kilobase exon per million mapped reads (FPKM); tags per million mapped reads; and counts) were generated using RSEM. Differential gene expression was calculated using DESeq2. To calculate FC cutoffs, datasets were normalized based on FPKM and purged of micro-RNAs, sno-RNAs and sca-RNAs. When multiple transcripts were detected for one gene, only the most abundant (highest average RPKM across all replicates) was analyzed. Transcripts with FPKM < 1 were excluded. Downstream analyses were performed with R 4.3.2, Morpheus (https://software.broadinstitute.org/morpheus/), VolcanosER (https://huygens.science.uva.nl/VolcaNoseR/), DAVID (https://davidbioinformatics.nih.gov), and Metascape (https://metascape.org/gp/index.html#/main/step1). Metascape enrichment scores were calculated using the formula *nN/kM*, where *N* = total genes in the study pool, i.e. human/mouse genome, *k* = genes in the pathway of interest, *M* = # genes in the input gene list, and n = overlap between *k* and M.

### Geneset Enrichment Analysis (GSEA)

For GSEA (https://www.gsea-msigdb.org/gsea/index.jsp) analysis (Mootha et al., 2003; Subramanian et al., 2005), enrichment score curves and member ranks were generated by GSEA software (Broad Institute). The Th9 cassette was obtained from a previously published analysis of Th9 cells from our group(Son et al., 2023). Other T helper subset cassettes were also obtained from previously published datasets from our group(Schwartz et al., 2019). Other pathways were analyzed based on the Human Molecular Signatures Database (MSigDB) collections.

### Single cell RNA sequencing analysis

Primary analysis of CITE-sequencing data was done using Seurat 5.2,0 (Satija et al., 2015). UMI files were downloaded for plaque-resident and circulating (pbmc) T cells (https://figshare.com/s/c00d88b1b25ef0c5c788). For all downstream analyses, UMI plots of plaque-resident and circulating CD4+ T cells were extracted by making a dataframe labeling each individual cell (column) based on CITE-seq antibody labeling of CD4^+^ and CD8+ T cells, then extracting a spreadsheet of samples that only included cells labeled as “CD4.Tcells.plaque” that were positive (by CITE-seq labeling) for CD4 but not for CD8. All downstream analysis was performed on the UMI matrix for the “CD4.Tcells.plaque” and “CD4.Tcells.pbmc” samples. UMAP plots were created and SPI1+ cells visualized using standard pipelines. To determine enrichment of the “allergic Th9” cassette, GSEA was performed on the “CD4.Tcells.plaque” and “CD4.Tcells.pbmc” UMI files, comparing cells in which *SPI1* expression was detected vs. cells in which *SPI1* expression was not detected, with “signal to noise” used to rank genes and “phenotype” permutation. For pathway analysis, Pearson correlations were calculated for *SPI1* in the “CD4.Tcells.plaque” file, and genes with a correlation score ≥ 0.2 were selected for pathway analysis.

### Western blotting

Endothelial cells were cultured as above and treated with either vehicle (1X PBS supplemented with 0.1% recombinant human serum albumin) or recombinant hIL-9 (PeproTech). Cells were washed twice with ice-cold PBS then lysed at the indicated time points in RIPA buffer containing protease & phosphatase inhibitor cocktail (Thermofisher, #78445). Cells were scraped and underwent microcentrifugation at 4**°**C and 14000g. The supernatant was collected for analysis and protein concentration was quantified using a Pierce BCA Protein Assay Kit (Thermofisher, # 23225). Equal protein concentrations were run on Mini-Protean TGXgel (BioRad) and transferred to nitrocellulose membranes by semi-dry transfer (Trans-blot turbo transfer system, BioRad). Nitrocellulose membranes were blocked for 1 hour at room temperature with agitation using 5% non-fat dry milk (NFDM) or 2% bovine serum albumin (BSA) for phospho-antibodies. Following the blocking step, membranes were incubated with primary antibody diluted 1:1000 in either 5% NFDM or BSA overnight at 4**°**C with agitation. Following incubation with primary antibody, membranes were washed with TBS supplemented with 0.1% Tween-20 detergent (TBS-T) three times for 10 minutes each with agitation. After the wash step, membranes were incubated for 1 hour at room temperature with either anti-rabbit or anti-mouse IgG linked to horseradish peroxidase (Cell Signaling Technologies) diluted 1:5000 in the appropriate blocking solution. Detailed information about antibodies can be found in **Supplementary Table 3**. After incubation with secondary antibody, the membranes were again washed in TBS-T 3 times 10 minutes each. Membranes were transferred to a clean and dry container and incubated for 5 minutes in SuperSignal West Pico Plus chemiluminescent substrate (ThermoFisher # 34580), followed by imaging on a Fluor Chem E imager (Bio Techne) or CCD imager (Azure Biosystems). All experiments used 3-6 biological replicates.

### Immunoprecipitation

Endothelial cells were cultured as above and lysed with 1X non-denaturing immunoprecipitation lysis buffer (Cell Signaling Technologies). Cell extracts were centrifuged at 4**°**C and 14000g for 10 minutes and sonicated if necessary. The supernatant was collected, and the cell pellet was discarded. Protein concentration was quantified using a Pierce BCA Protein Assay Kit (ThermoFisher). 1 milligram of total input protein from the cell lysate was incubated with IL9R polyclonal antibody (Biolegend) or isotype IgG at a 1:20 dilution overnight in protein low-bind tubes with rotation at 4**°**C. Following the overnight incubation step, 40 uL of Protein A/G magnetic bead slurry (ThermoFisher) per sample was mixed with 460 uL of wash buffer and gently vortexed to mix. The magnetic beads were then precipitated with a magnetic stand and washed once more with buffer. Following the bead wash steps, the cleaned magnetic beads were incubated with the antibody-cell lysate mixture overnight at 4**°**C with rotation. Following this incubation step, the beads were collected with a magnetic stand and washed twice with binding and wash buffer followed by a final wash with deionized water. After this final wash step, the beads were treated with 50 uL of elution buffer (0.1 M glycine, pH 2.0) and mixed with 4X loading buffer for Western blot (BioRad). The final elution was then analyzed via Western blot as described above. IL-2R γ rabbit monoclonal antibody (Cell Signaling Technologies) was used to detect co-immunoprecipitated IL-2R as above.

### Statistics

For all experiments other than transcriptome-wide analysis, statistics were calculated in GraphPad (Prism). Samples were analyzed to determine whether distribution was normal or non-normal. Normally distributed samples were analyzed using paired or unpaired t-test, depending on experimental design. Non-normally distributed samples were analyzed using Wilcoxon rank sum test (paired) or Mann-Whitney U test (unpaired). Significance between groups was determined using ANOVA. For experiments using multiple comparisons, multiple comparison adjustment was performed using Benjamini-Hochberg correction. All statistical testing was two-sided.

## Supporting information

Supplementary Tables (all)

Supplementary Figures (all)

## Author Contributions

IB performed experiments, analyzed data, and wrote the manuscript. YB designed the study, performed and supervised experiments, analyzed data, and wrote the manuscript. AB, MS, MMK, GH, CAGH, BT, AS, KC, JK, and KJ performed experiments. AD, JR, and NNM provided patient samples and data, and performed statistical analysis. JDP performed histological analysis. MC provided representative coronary CT images. PAFG, JDM, TMPW, and NNM developed study protocols, provided patient samples and data, and edited the manuscript. LG, AF, AG, and TS analyzed data from human cadaveric coronary artery specimens. GHF designed the study, performed and supervised experiments, analyzed data, and wrote the manuscript. DMS designed the study, performed and supervised experiments, analyzed data, provided funding, supervised the study, and wrote the manuscript.

## Acknowledgments

The authors would like to thank the NHLBI Microscopy core (Dr. Christian Combs), the NHLBI Histology core (Dr. Zu-Xi Yu), Dr. Laurel G. Mendelsohn, and Dr. Dr. Abhinav Saurabh.

---

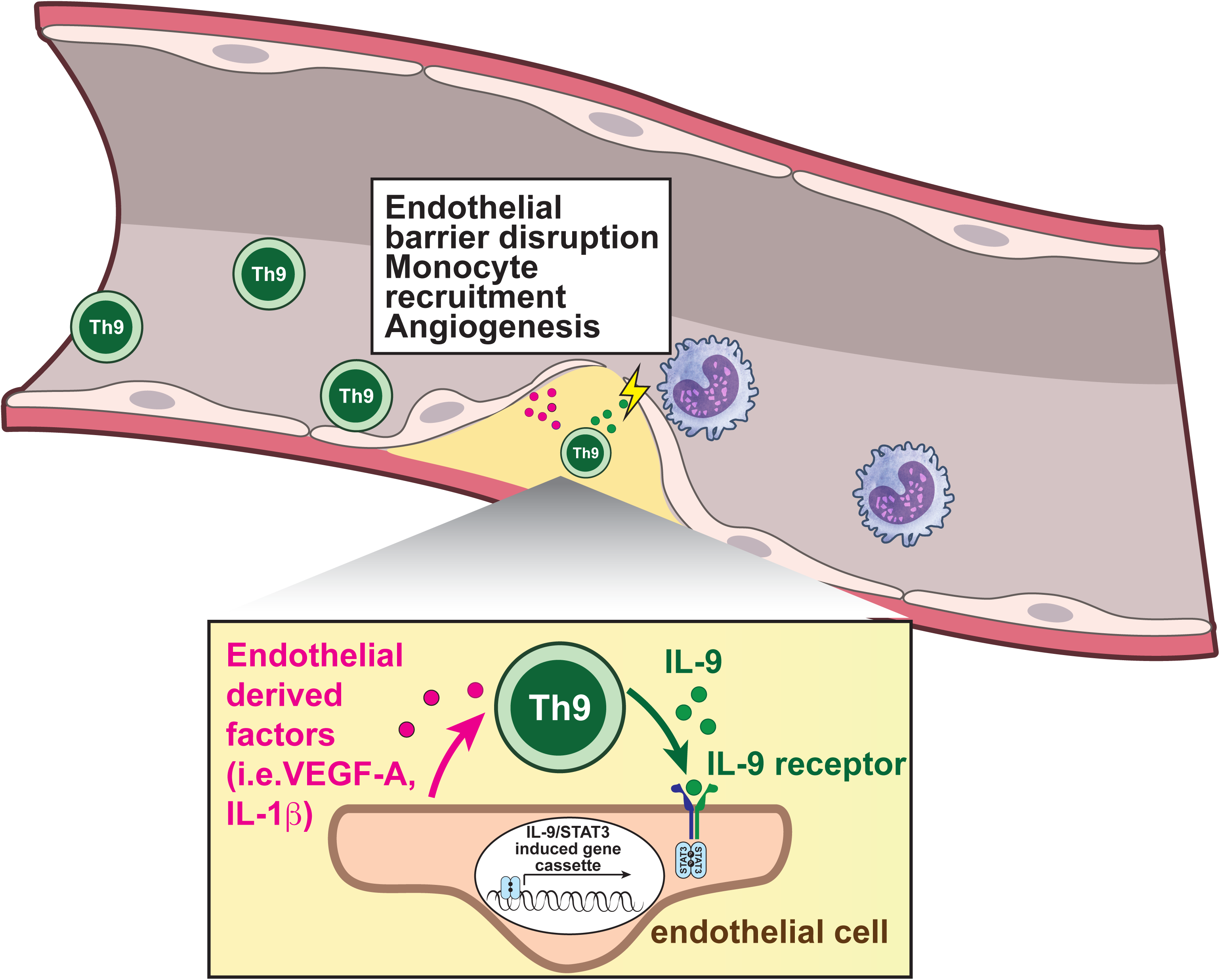

