## Supplementary Tables (all) for "Th9-endothelial cell crosstalk promotes inflammatory atherosclerotic cardiovascular disease"

**Supplementary Table 1. Characterization of Th9^high^ vs. Th9^low^ Psoriasis Patients at Baseline**

| **Variable** | **Th9^low^** (<0.90%) | **Th9^high^** (≥0.90%) | **P-value** |
| --- | --- | --- | --- |
| **Demographic and Clinical Characteristics** | n=12 | n=8 |  |
| Age, years | 48.8 ± 13.0 | 53.5 ± 12.0 | 0.42 |
| Males, N (%) | 6 (50) | 7 (88) | 0.16 |
| Hypertension, N (%) | 3 (27) | 4 (50) | 0.31 |
| Hyperlipidemia, N (%) | 4 (33) | 4 (50) | 0.46 |
| Type-2 diabetes mellitus, N (%) | 2 (17) | 2 (25) | 0.65 |
| Statin therapy, N (%) | 0 (0) | 4 (50) | **0.01** |
| Current Smoker, N (%) | 0 (0) | 1 (13) | 0.21 |
| Body Mass Index, kg/m2 | 27.7 (26.7-31.7) | 29.0 (25.9-31.7) | 0.73 |
| Waist to Hip Ratio | 0.95 (0.90-1.05) | 1.0 (0.97-1.02) | 0.31 |
| Framingham Risk Score | 1.11 (0.45-3.60) | 5.42 (2.46-8.38) | 0.06 |
| Psoriasis Area Severity Index Score | 2.8 (2.5-5.5) | 3.5 (3.2-8.2) | 0.19 |
| Biologic Therapy, N (%) | 1 (8) | 0 (0) | 1.00 |
| **Clinical and Lab Values** |  |  |  |
| Total Cholesterol, mg/dL | 208 (183-233) | 167 (158-171.5) | **0.003** |
| HDL Cholesterol, mg/dL | 60 (45-70) | 50 (38.5-67) | 0.49 |
| LDL Cholesterol, mg/dL | 114.5 (106-142) | 86.5 (80-94.5) | **0.003** |
| Triglycerides, mg/dL | 105 (67-139) | 129.5 (63-174.5) | 0.67 |
| Cholesterol Efflux Capacity, % | 1.02 ± 0.14 | 1.07 ± 0.10 | 0.4 |
| High-sensitivity C-reactive protein, mg/dL | 1.6 (1.1-3.5) | 2.2 (1.0-4.1) | 0.55 |
| IL-9 (%of CD4^+^CD45RO^+^ T cells) | 0.34 (0.28-0.50) | 2.11 (1.05-3.61) | **<0.001** |
| Systolic Blood Pressure, mmHg | 122.8 ± 10.3 | 114.4 ± 8.7 | 0.07 |
| Diastolic Blood Pressure, mmHg | 72.7 ± 6.0 | 70.1 ± 11.8 | 0.53 |
| **Coronary Characteristics** |  |  |  |
| Non-Calcified Burden, % | 0.98 (0.77-1.24) | 1.37 (0.98-1.64) | **0.004** |
| Total Burden, % | 1.01 (0.77-1.24) | 1.41 (1.17-2.08) | **0.001** |

Results are shown for participants from study 13H-0065 for whom T helper phenotypes and coronary computed tomography data (n = 20). Continuous variables are expressed as mean ± standard deviation and categorical variables as total N (%).Normal continuous variables reported as mean ± S.D., non-normal continuous variables reported as med (IQR) and categorical variables reported as N (%).

**Supplementary Table 2.Other cytokines in CD45RO+ CD4+ T cells; Non-Calcified Burden Adjusted Regression Analysis**

| **Variable (cytokine, % positive of CD4^+^ CD45RO^+^ T cells)** | **Beta** | **P-Value*** |
| --- | --- | --- |
| Unadjusted (IFN-γ) | 0.24 | 0.08 |
| Adjusted for BMI, FRS, and Statin Use (IFN-γ) | 0.06 | 0.42 |
| Unadjusted (IL-4) | 0.14 | 0.31 |
| Adjusted for BMI, FRS, and Statin Use (IL-4) | 0.02 | 0.78 |
| Unadjusted (IL-13) | 0.01 | 0.97 |
| Adjusted for BMI, FRS, and Statin Use (IL-13) | -0.05 | 0.49 |
| Unadjusted (IL-17A) | -0.31 | **0.02** |
| Adjusted for BMI, FRS, and Statin Use (IL-17A) | 0.14 | 0.12 |

*Multivariable regression analysis between non-calcified burden and % cytokine-positive CD45RO+ CD4+ T cells unadjusted or adjusted for BMI, FRS, and statin use.

**Supplementary Table 3: Antibodies used for flow cytometry, Western blotting, immunofluorescent staining**

| **Antibody** | **Catalog number** | **Concentration** |
| --- | --- | --- |
| Anti-mouse F4/80- SB 600 | Invitrogen, # 63480182 | 1:200 |
| Anti-mouse/human CD11b- FITC | BioLegend, # 101206 | 1:200 |
| Anti-mouse SiglecF-PE | BD Pharmingen, # 552126 | 1:200 |
| Anti-mouse Ly6G- PerCP/Cy5.5 | BioLegend, # 127616 | 1:200 |
| Anti-mouse Ly6C- Alexa Fluor 647 | BioLegend, # 128010 | 1:200 |
| Anti-mouse Ly6C- Alexa Fluor 700 | BioLegend, # 128024 | 1:200 |
| Anti-mouse CD3- Brilliant Violet 421 | BioLegend, # 100336 | 1:200 |
| Anti-mouse CD45.2- PE-Cy7 | BioLegend, # 109830, | 1:200 |
| anti-mouse CD45- APC-Cy7 | BioLegend, # 157618 | 1:200 |
| Anti-mouse TCRβ- APC-Cy7 | BioLegend, # 109220 | 1:200 |
| Anti-mouse TCRβ- SB 645 | Invitrogen, # 64596182 | 1:200 |
| Anti-mouse CD4- PE/Dazzle 594 | BioLegend, # 100566 | 1:200 |
| Anti-mouse CD4- PerCP-Cy5.5 | BioLegend, # 100540 | 1:200 |
| Anti-mouse CD8a- Alexa Fluor 700 | BioLegend, # 100730 | 1:200 |
| Anti-mouse IL9- PE | BioLegend, # 514104 | 1:100 |
| Anti-mouse IL13- PE-Cy7 | Invitrogen, # 25713382 | 1:200 |
| Anti-mouse IL17- APC | Invitrogen, # 17717781 | 1:200 |
| Anti-mouse IFNγ- Pacific Blue | BioLegend, # 505818 | 1:200 |
| anti-mouse IFNγ-PE | BioLegend, #505808 | 1:200 |
| Anti-mouse FOXP3- eFlr 450 | Invitrogen, # 48577382 | 1:200 |
| Anti-mouse CD19- APC | Invitrogen, # 17019380 | 1:200 |
| Anti-mouse B220- PE | Pharmingen, # 01125B | 1:200 |
| Anti-mouse IL9R- PE-Cy7 | BioLegend, # 158807 | 1:100 |
| Anti-human CD3, APC-Cy7 | Biolegend, # 344817 | 1:200 |
| Anti-human CD3, AF700 | BD Bioscience, # 557943 | 1:200 |
| Anti-human CD4, BUV395 | BD bioscience, # 564724 | 1:200 |
| Anti-human CD4, APC | Invitrogen, # MHCD0405 | 1:200 |
| Anti-human CD8, APC-Cy7 | BD Bioscience, # 557760 | 1:200 |
| Anti-human CD14, FITC | BD Bioscience, 555397 | 1:200 |
| Anti-human CD16, FITC | Biolegend, 302006 | 1:200 |
| Anti-human CD19, AF488 | Biolegend, 302206 | 1:200 |
| Anti-human CD45RO, PE-Texas Red | Beckmann Coulter, #IM2712U | 1:200 |
| Anti-human CD45RA, BV711 | BD Bioscience, # 563733 | 1:200 |
| Anti-human IL-2, PerCpCy5.5 | Biolegend, # 500322 | 1:200 |
| Anti-human IL-4, FITC | Biolegend, # 500806 | 1:200 |
| Anti-human IL-4, BV605 | Biolegend, # 500828 | 1:200 |
| Anti-human IL-5, BV421 | Biolegend, # 504311 | 1:200 |
| Anti-human IL-9, MH9A4 | Biolegend, # 507605 | 1:200 |
| Anti-human IL-10, PE-Cy7 | Biolegend, # 501420 | 1:200 |
| Anti-human IL-13, BV711 | BD Bioscience, # 564288 | 1:200 |
| Anti-human IL-17A, APC | invitrogen, # 50-7178-42 | 1:200 |
| Anti-human IL-17A, AF700 | BD Bioscience, # 557995 | 1:200 |
| Anti-human IL-17A, v450 | BD Bioscience, # 560371 | 1:200 |
| Anti-human IFN-γ, AF700 | BD Bioscience, # 557995 | 1:200 |
| Anti-human IFN-γ, v450 | BD Bioscience, # 560371 | 1:200 |
| Rabbit anti-human PU.1 | Cell Signaling Technology, # 2258 | 1:400 |
| Mouse anti-human IL9R | ThermoFisher, # MA5-28556 | 1:400 (IHC) |
| Mouse anti-human IL9R | Biolegend, #1653 | 1:20 (IP); 1:400 (IHC), 1:1000 (WB) |
| Rabbit anti-human IL2RG | Cell Signaling Technology, # 49622 | 1:1000 |
| Phalloidin-g, AF488 | ThermoFisher, # A12379 | 1:500 |
| Goat anti-mouse/human VE-Cadherin | R&D, # AF1002 | 1:100 |
| Mouse anti-mouse/human VE-Cadherin | Santa Cruz, # sc-9989 | 1:100 |
| Rabbit anti-human CD3 | Roche, # 10243295001 | 1:200 |
| Rabbit anti-human CD4 | Roche, # 05552737001 | 1:200 |
| Rabbit anti-mouse/human pSTAT3 (Y705) | Cell Signaling Technology, # 9145 | 1:1000 |
| Rabbit anti- mouse/human pSTAT1 (Y701) | Cell Signaling Technology, # 9167 | 1:1000 |
| Mouse anti- mouse/human STAT3 | Cell Signaling Technology, # 9139 | 1:1000 |
| Rabbit anti- mouse/human STAT1 | Cell Signaling Technology, # 14994 | 1:1000 |
| Rabbit anti-human beta actin | Cell Signaling Technology, # 4970 | 1:1000 |
| AF555 anti-rabbit IgG | ThermoFisher, # A31572 | 1:150 |
| AF488 anti-mouse IgG | ThermoFisher, # A21202 | 1:150 |
| DAPI | ThermoFisher, # D1306 | 1:500 |
| AF488 anti-goat IgG | ThermoFisher, # A11055 | 1:150 |
| AF555 anti-rabbit IgG | ThermoFisher, # A31572 | 1:150 |
| AF555 anti-mouse IgG | ThermoFisher, # A31570 | 1:150 |
