## Supplementary Figures (all) for "Th9-endothelial cell crosstalk promotes inflammatory atherosclerotic cardiovascular disease"

### Supplementary Figure 1

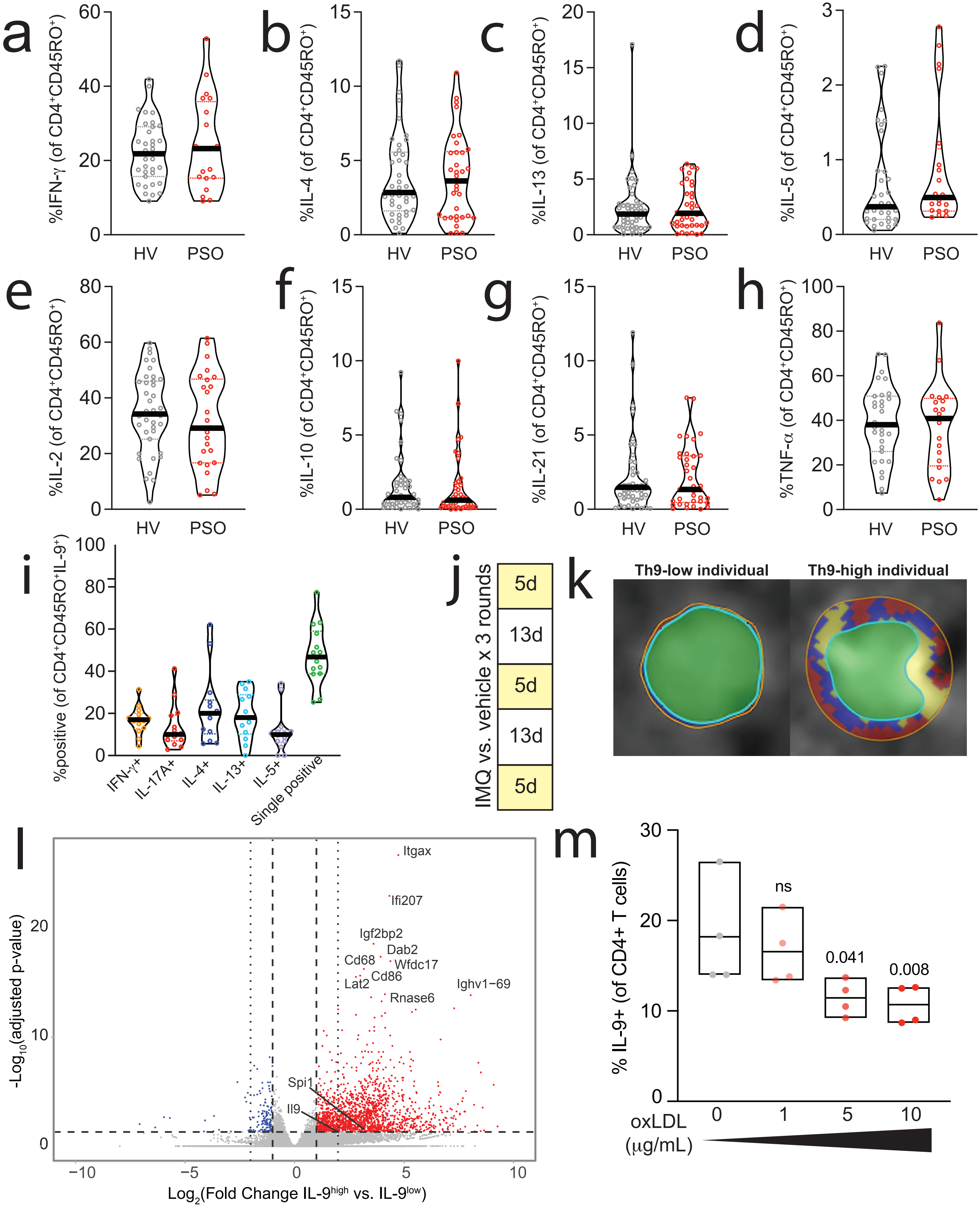

Supplementary Figure 2

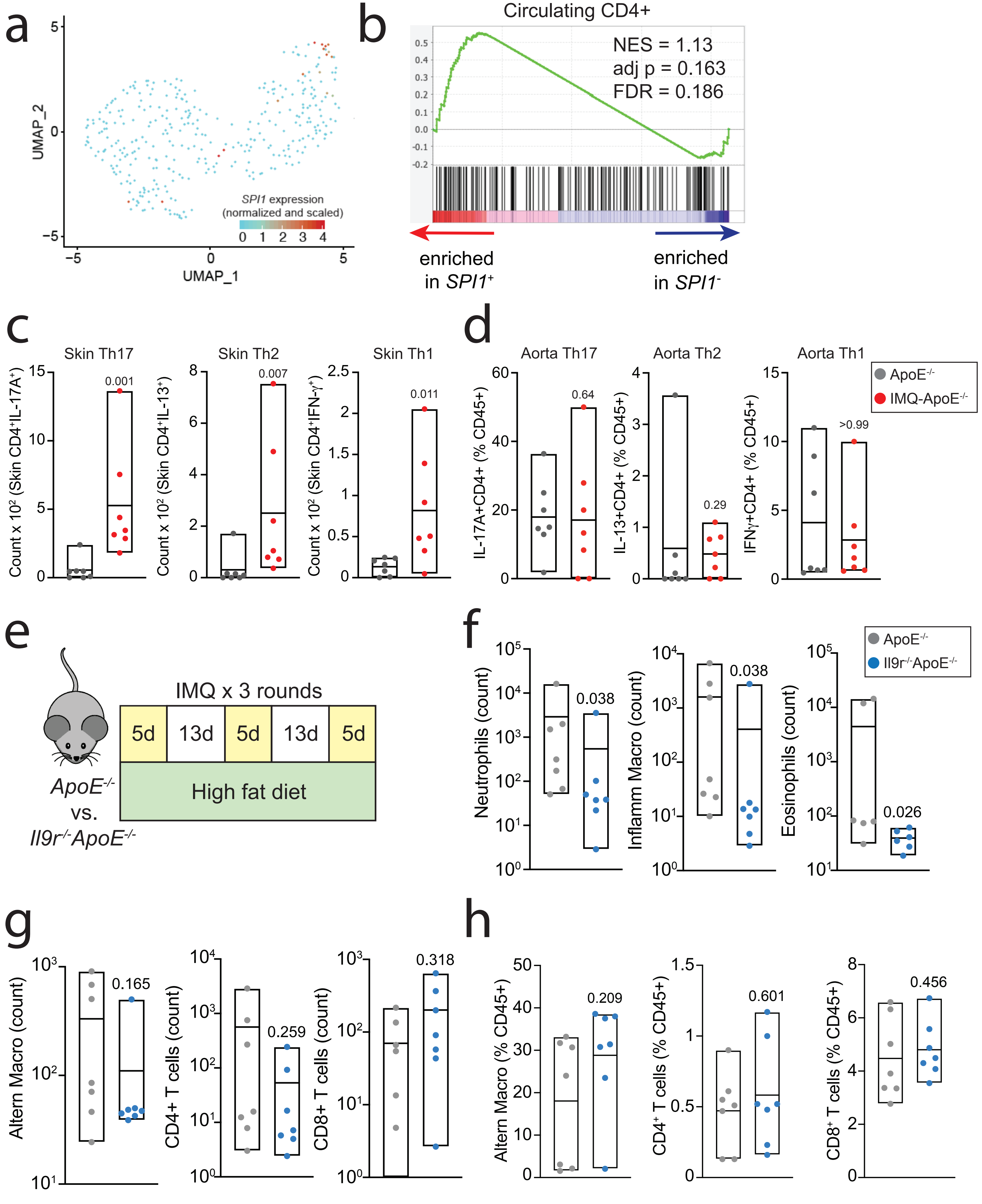

### Supplementary Figure 3

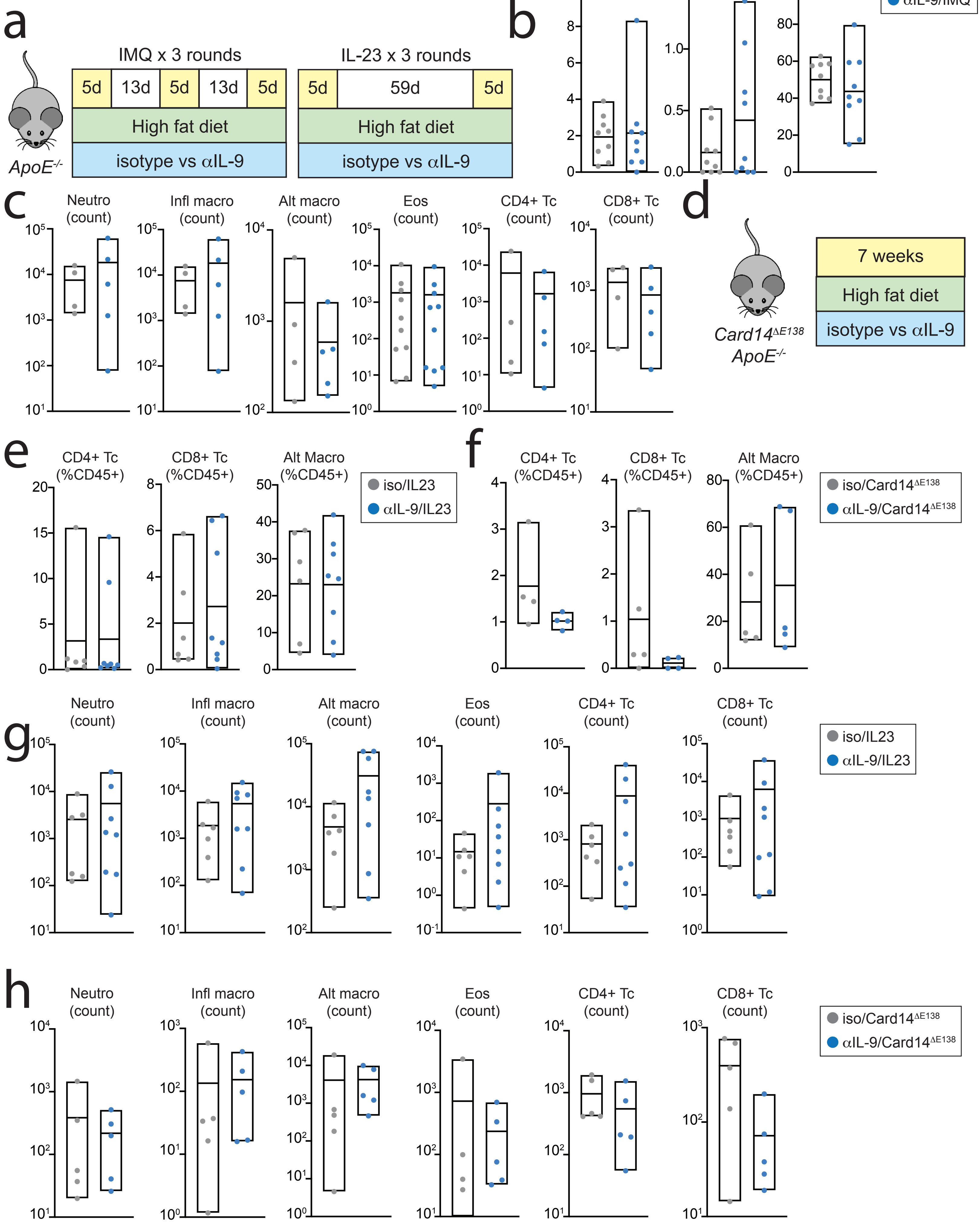

### Supplementary Figure 4

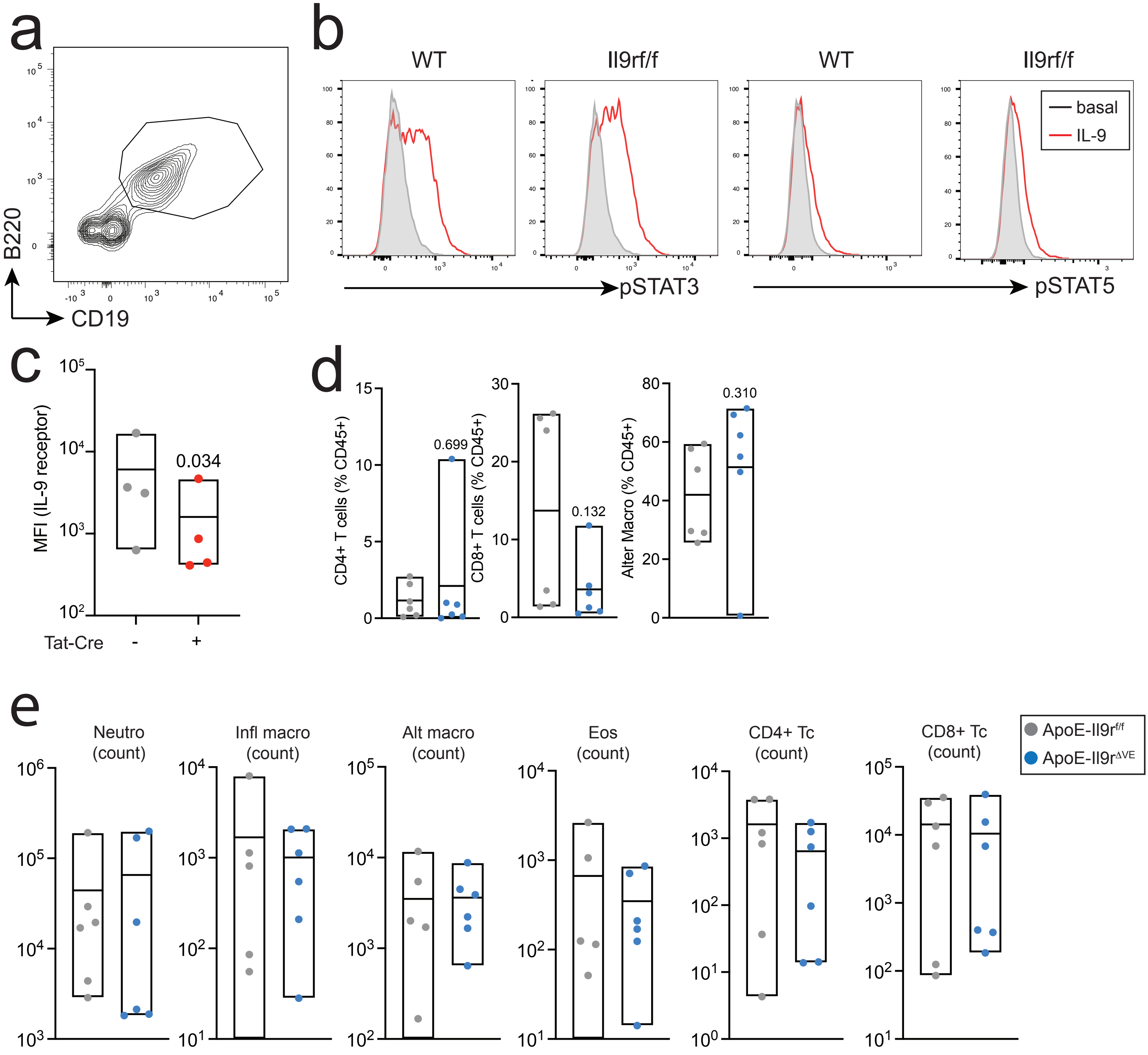

### Supplementary Figure 5

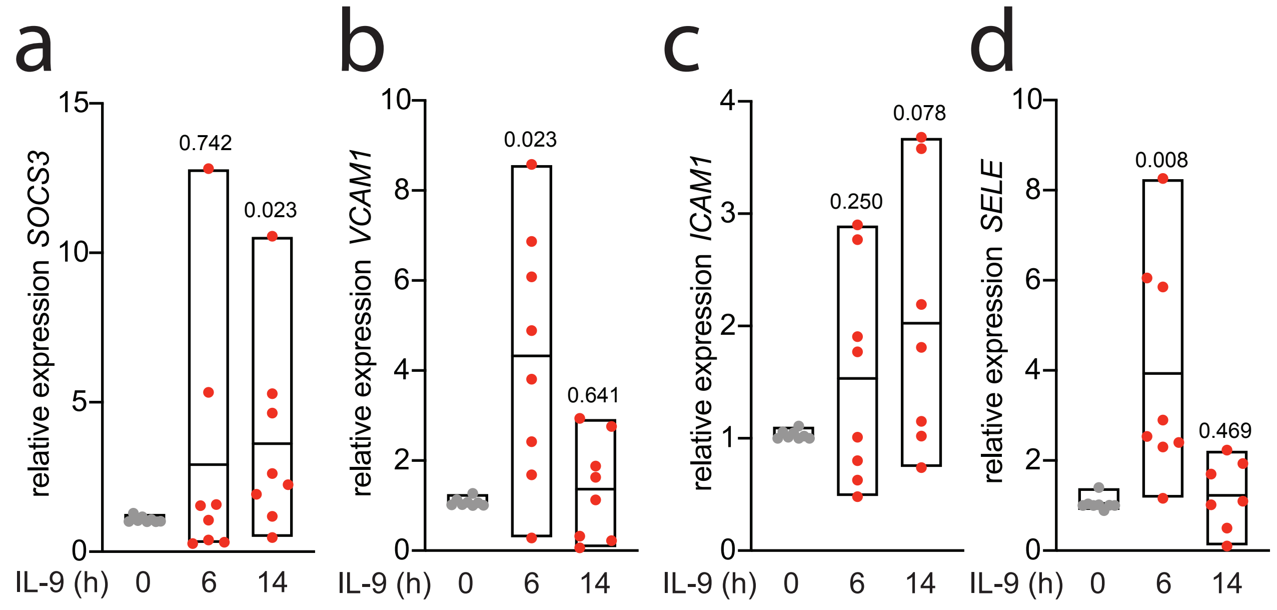
