## Supplementary Figure Legends for "Th9-endothelial cell crosstalk promotes inflammatory atherosclerotic cardiovascular disease"

**Supplementary Figure 1.** a-h. Violin plots show % circulating T cells (CD45+/CD3+/CD4+) positive for IFN-γ (a), IL-4 (b), IL-13 (c), IL-5 (d), IL-2 (e), IL-10 (f), IL-21 (g), and TNF-α (h) cells in non-psoriatic healthy volunteers (HV, n = 47, grey) and patients with psoriasis (PSO, n = 36, red). i. Violin plot shows % circulating Th9 cells (CD45+/CD3+/CD4+/IL-9+) also staining positive for hallmark effector cytokines for Th1 (IFN-γ), Th17 (IL-17A), Th2 (IL-4, IL-13, IL-5), or not staining positive for any other hallmark T effector cytokine (single positive). j. Schematic shows chronic imiquimod (IMQ) induced skin inflammation model in INFER (IL-9) reporter mice. k. Representative image shows calcified and non-calcified coronary plaque in a subject with psoriasis and an age/sex matched non-psoriatic volunteer. Red, low attenuation; Blue, fibro fatty plaque; yellow,= calcified plaque; green, lumen; aqua, inner vessel wall or intima layer; orange, outer vessel wall or adventitia. l. Volcano plot shows genes differentially expressed in Th9 (CD45+TCRβ+CD4+CD8-GFP+) vs. non-Th9 (CD45+TCRβ+CD4+CD8-GFP-) cells sorted from INFER mice treated with IMQ as in S1j.

**Supplementary Figure 2.** a. UMAP plot shows plaque-resident CD4+ T cells from CITE-seq data of plaque-resident CD4+ T cells from humans undergoing carotid endarterectomy (<https://figshare.com/s/c00d88b1b25ef0c5c788>). Normalized and scaled *SPI1* expression is shown, including SPI1-negative cells (blue) and SPI1-positive cells (red). b. GSEA plot shows enrichment of “allergic Th9 cassette” in circulating resident *SPI1*+ (Th9) vs. *SPI1*- (non-Th9) cells from the same patient population in Fig S2a. c. Bar graphs show counts of skin-infiltrating Th17 (CD45+TCRβ+CD4+CD8-IL-17A+), Th2 (CD45+TCRβ+CD4+CD8-IL-13+), and Th1 (CD45+TCRβ+CD4+CD8-IFN-γ+) cells, normalized to skin weight. d. Bar graphs show percent Th17 (TCRβ+CD4+CD8-IL-17A+), Th2 (TCRβ+CD4+CD8-IL-13+), and Th1 (TCRβ+CD4+CD8-IFN-γ+) of aorta-infiltrating CD45+ cells. e. Schematic shows chronic imiquimod (IMQ) induced inflammatory atherogenesis model in *Il9r^+/+^ApoE^-/-^*and *Il9r^-/-^ApoE^-/-^* mice. f,g. Bar graphs show counts of skin-infiltrating neutrophils (CD45+CD11c+CD3-Ly6G+, f), eosinophils (CD45+CD11c+CD3-SiglecF+F4/80-, f), inflammatory macrophages (CD45+CD11c+CD3-SiglecF+F4/80+CD11b+, f), alternative macrophages (CD45+CD11c+CD3-SiglecF+F4/80+CD11b-, g), CD4+ T cells (CD45+CD11c-CD3+TCRβ+CD4+CD8-, g), and CD8+ T cells (CD45+CD11c-CD3+TCRβ+CD4-CD8+,g), normalized to skin weight. h. Bar graphs show percent alternative macrophages, CD4+ T cells, and CD8+ T cells, of aorta-infiltrating CD45+ cells. For all graphs, statistics by Mann-Whitney.

**Supplementary Figure 3.** a. Schematic shows chronic imiquimod (IMQ) induced inflammatory atherogenesis model and IL-23 induced inflammatory atherogenesis model in *ApoE^-/-^* mice treated with IL-9-blocking antibody or isotype control. b. Bar graphs show percent CD4+ T cells (CD45+CD11c-CD3+TCRβ+CD4+CD8-), CD8+ T cells (CD45+CD11c-CD3+TCRβ+CD4-CD8+), and alternative macrophages (CD45+CD11c+CD3-SiglecF+F4/80+CD11b-) of aorta-infiltrating CD45+ cells in mice exposed to the chronic IMQ induced inflammatory atherogenesis model and treated with IL-9-blocking antibody vs. control. c. Bar graphs show counts of skin-infiltrating neutrophils (CD45+CD11c+CD3-Ly6G+), inflammatory macrophages (CD45+CD11c+CD3-SiglecF+F4/80+CD11b+), alternative macrophages (CD45+CD11c+CD3-SiglecF+F4/80+CD11b-), eosinophils (CD45+CD11c+CD3-SiglecF+F4/80-),CD4+ T cells (CD45+CD11c-CD3+TCRβ+CD4+CD8-), and CD8+ T cells (CD45+CD11c-CD3+TCRβ+CD4-CD8+), normalized to skin weight in mice exposed to the chronic (IMQ) induced inflammatory atherogenesis model and treated with IL-9-blocking antibody vs. control. d. Schematic shows *Card14^ΔE138^ApoE^-/-^* atherogenesis model in mice treated with IL-9-blocking antibody or isotype control e,f. Bar graphs show percent CD4+ T cells (CD45+CD11c-CD3+TCRβ+CD4+CD8-), CD8+ T cells (CD45+CD11c-CD3+TCRβ+CD4-CD8+), and alternative macrophages (CD45+CD11c+CD3-SiglecF+F4/80+CD11b SiglecF+F4/80+CD11b-) of aorta-infiltrating CD45+ cells in mice exposed to the IL-23 induced inflammatory atherogenesis model (e) and *Card14^ΔE138^ApoE^-/-^* atherogenesis model (f) and treated with IL-9-blocking antibody vs. control. g,h. Bar graphs show counts of skin-infiltrating neutrophils (CD45+CD11c+CD3-Ly6G+), inflammatory macrophages (CD45+CD11c+CD3-SiglecF+F4/80+CD11b+), alternative macrophages (CD45+CD11c+CD3-SiglecF+F4/80+CD11b-), eosinophils (CD45+CD11c+CD3-SiglecF+F4/80), CD4+ T cells (CD45+CD11c-CD3+TCRβ+CD4+CD8-), and CD8+ T cells (CD45+CD11c-CD3+TCRβ+CD4-CD8+), normalized to skin weight in mice exposed to the IL-23 induced inflammatory atherogenesis model (g) and *Card14^ΔE138^ApoE^-/-^* atherogenesis model (h) and treated with IL-9-blocking antibody vs. control.

**Supplementary Figure 4.** a. Representative flow plot shows *in vitro* differentiated memory B cells based on CD19 and B220 expression. b. Representative histograms show basal (grey) and IL-9-induced (red) STAT3 and STAT5 phosphorylation in memory B cells from WT and *Il9r^f/f^* mice. c. Pooled mean fluorescence intensity (flow cytometry) for IL-9 receptor staining in *Il9r^f^*^/f^ B cells treated with vehicle vs Tat-Cre *in vitro.* d. Bar graphs show percent CD4+ T cells (CD45+CD11c-CD3+TCRβ+CD4+CD8-), CD8+ T cells (CD45+CD11c-CD3+TCRβ+CD4-CD8+), and alternative macrophages (CD45+CD11c+CD3-SiglecF+F4/80+CD11b SiglecF+F4/80+CD11b-) of aorta-infiltrating CD45+ cells in *Il9r^f/f^ApoE^-/-^*and *Il9r^ΔVE^ApoE^-/-^* mice exposed to the IMQ induced inflammatory atherogenesis model. e. Bar graphs show counts of skin-infiltrating neutrophils (CD45+CD11c+CD3-Ly6G+), inflammatory macrophages (CD45+CD11c+CD3-SiglecF+F4/80+CD11b+), alternative macrophages (CD45+CD11c+CD3-SiglecF+F4/80+CD11b-), eosinophils (CD45+CD11c+CD3-SiglecF+F4/80), CD4+ T cells (CD45+CD11c-CD3+TCRβ+CD4+CD8-), and CD8+ T cells (CD45+CD11c-CD3+TCRβ+CD4-CD8+), normalized to skin weight in *Il9r^f/f^ApoE^-/-^*and *Il9r^ΔVE^ApoE^-/-^* mice exposed to the IMQ induced inflammatory atherogenesis model.

**Supplementary Figure 5.** a-d Bar graphs show gene expression (normalized to *TBP*) for *SOCS3* (a), *VCAM1* (b), *ICAM1* (c), and *SELE* (d) in HAoEC treated with vehicle vs. IL-9 (n = 8). For d-h, p-values, Wilcoxon.
